# Next generation cytogenetics: comprehensive assessment of 48 leukemia genomes by genome imaging

**DOI:** 10.1101/2020.02.06.935742

**Authors:** Kornelia Neveling, Tuomo Mantere, Susan Vermeulen, Michiel Oorsprong, Ronald van Beek, Ellen Kater-Baats, Marc Pauper, Guillaume van der Zande, Dominique Smeets, Daniel Olde Weghuis, Marian J Stevens-Kroef, Alexander Hoischen

**Author notes:** Correspondence: Alexander Hoischen, PhD, Department of Human Genetics & Department of Internal Medicine, Radboud university medical center, 6500HB Nijmegen, The Netherlands. These authors contributed equally to this work (Kornelia Neveling, Tuomo Mantere).

## Abstract

Somatic structural variants are important for cancer development and progression. In a diagnostic set-up, especially for hematological malignancies, the comprehensive analysis of all cytogenetic aberrations in a given sample still requires a combination of techniques, such as karyotyping, fluorescence *in situ* hybridization and CNV-microarrays. We hypothesize that the combination of these classical approaches could be replaced by high-resolution genome imaging.

Bone marrow aspirates or blood samples derived from 48 patients with leukemia, who received a clinical diagnoses of different types of hematological malignancies, were processed for genome imaging with the Bionano Genomics Saphyr system. In all cases cytogenetic abnormalities had previously been identified using standard of care workflows. Based on these diagnostic results, the samples were divided into two categories: simple cases (<5 aberrations, n=37) and complex cases (≥5 aberrations or an unspecified marker chromosome, n=11). By imaging the labelled ultra-long gDNA molecules (average N50 >250kb), we generated on average ∼280-fold mapped genome coverage per sample. Chromosomal aberrations were called by Bionano Genomics Rare variant pipeline (RVP) specialized for the detections of somatic variants.

Per sample, on average a total of 1,454 high confidence SVs were called, and on average 44 (range: 14-130) of those were rare *i.e*. not present in the population control database. Importantly, for the simple cases, all clinically reported aberrations with variant allele frequencies higher than 10% were detected by genome imaging. This held true for deletions, insertions, inversions, aneuploidies and translocations. The results for the complex cases were also largely concordant between the standard of care workflow and optical mapping, and in several cases, optical mapping revealed higher complexity than previously known. SV and CNV calls detected by optical mapping were more complete than any other previous single test and likely delivered the most accurate and complete underlying genomic architecture. Even complex chromothripsis structures were resolved. Finally, optical mapping also identified multiple novel events, including balanced translocations that lead to potential novel fusion-genes, opening the potential to discover new prognostic and diagnostic biomarkers.

The full concordance with diagnostic standard assays for simple cases and the overall great concordance with (previously likely incompletely understood) complex cases demonstrates the potential to replace classical cytogenetic tests with genome imaging. In addition, this holds the potential to rapidly map new fusion genes and identify novel SVs and CNVs as novel potential leukemia drivers.

## Introduction

The introduction of next generation sequencing (NGS) has dramatically changed the way clinical molecular laboratories analyze their samples over the past 10 years. Sanger sequencing is rapidly losing ground compared to NGS, and single gene analyses are gradually replaced by gene panels, exomes and genomes.^1^ In clinical cytogenetics, a trend towards NGS-based analysis is visible since the introduction of non-invasive prenatal testing (NIPT)^2^ and other sequencing tests using cell-free DNA,^3^ but for most clinical cytogenetic analyses (a combination of) karyotyping, fluorescence *in situ* hybridization (FISH) and CNV-microarrays are still performed to detect genetic biomarkers of disease. Each of these tests has its own limitations; e.g. karyotyping has a maximum banding resolution of ∼5Mb; FISH has a higher resolution, but requires *a priori* knowledge of which loci to test and is limited in throughput; and CNV-microarrays offer the best resolution down to few kb, but lack the ability to identify balanced chromosomal aberrations including translocations, inversions. CNV-microarrays are also unable to map gained material; i.e. they cannot distinguish tandem duplications from insertions in trans.

In tumor genetics, and especially for hematological disorders, the choice for the respective diagnostic test depends on the underlying clinical diagnosis in combination with available and suitable tissues that can be tested. For several types of malignancies, the high degree of balanced translocations, some of which lead to cancer driving fusion-genes, still requires karyotyping and FISH as routine diagnostic assays. Different clinical testing guidelines define when to use which test in different political and geographical regions.^4; 5^ In our laboratory we use a combination of karyotyping and FISH for CML and lymphoma; karyotyping, FISH and CNV-microarray for AML and ALL; karyotyping and CNV-microarray for MDS and MPN; CNV-microarray for CLL; and FISH and CNV-microarray on CD138 enriched plasma cells for MM. Also of note, FISH represents multiple distinct tests, targeting different loci, that also vary for different clinical indications. At present, such divergence in diagnostic tests are accepted and seem unavoidable.

Here, we aimed to investigate whether clinical cytogenetics could become more generic by introducing a single test for cytogenetic assessment of leukemia samples: high-resolution genome imaging.

Genome-imaging of extremely long linear molecules, combined with optical mapping to detect SVs and CNVs, is an emerging technology that may have potential to replace all three above mentioned assays in cytogenetic diagnostic laboratories.^6-8^ Originally developed by Dr. David C. Schwartz and his lab at NYU in the 1990s,^9^ more recently genome imaging has been implemented in nanochannels arrays where high-throughput imaging of long, single DNA molecules (0.15-2.5 Mb) containing fluorescent labels marking sequence specific motifs distributed throughout the genome is achieved. Optical mapping is then able to reconstruct the genome, with highly accurate structure and contiguity in consensus maps up to chromosome arm length. Label pattern differences relative to a reference are detected and these differences are used to call structural variants (SVs).^10^ Because of the unique value gained by optical mapping of ultra-long DNA reads, it has been used in essentially all modern reference genome assemblies (human GRCh,^11; 12^ mouse,^13^ goat,^14^ maize, ^15^ as well as benchmark structural variation papers.^16-18^

The latest iteration of this technology, now marketed as genome imaging on the Bionano Genomics’ Saphyr system (Bionano Genomics, San Diego) generates images of molecules with average N50 >250kb and can generate ∼300x genome coverage per flow cell (3 flow cells per chip, 2 chips per instrument run). The ultra-long high molecular weight (UHMW) DNA molecules are fluorescently labelled on a 6-mer ssDNA motif (currently DLE1: CTTAGG), with an average label density of 15 labels per 100kb. Accurate and precise patterns of labels allows to a) *de novo* assemble the human genome which is then compared to the reference genome map; or b) extract aberrant molecules from reference alignments followed by local consensus generation, in order to detect SVs such as deletions, insertion, inversions, duplications, translocations, as well as copy number variants (CNVs) and whole chromosome aneuploidies in a genome-wide manner. The current technology allows SV detection down to 500bp resolution (for insertions and deletions, when using the *de novo* assembly pipeline), which is much higher compared to karyotyping, FISH and CNV-microarrays. Although the current methodology allows detection of balanced and unbalanced events, smaller insertions will have unknown origin when the inserted sequence is too small to contain a unique motif pattern, and breakpoint accuracy has median uncertainty of 3.1kb.^19^ It is also expected that balanced SVs with centromeric breakpoints will escape detection, and copy number neutral loss of heterozygosity (CN-LOH) detection is currently not enabled.

Here we describe a clinical validation study to investigate 48 leukemia samples with simple and complex cytogenetic aberrations using Bionano genome imaging. All 48 samples have been previously analyzed using karyotyping, FISH and/or CNV-microarray as part of routine diagnostic testing.

## Material and Methods

### Sample selection

Heparinized bone marrow or peripheral blood samples were sent to our clinical laboratory for routine cytogenetic diagnostic testing (karyotyping, FISH and/or genome wide CNV-microarray). In cases with sufficient left-over material, DNA stabilization buffer (Bionano Genomics) was added to heparinized bone marrow or blood and stored at -80 °C. Fifty samples with a cytogenetically abnormal result were anonymized and processed for Bionano optical mapping.

### Isolation of ultra-high molecular weight (UHMW) gDNA for genome imaging

UHMW gDNA was isolated from heparinized bone marrow aspirates (BMA) and EDTA- or heparin-blood stored at -80°C following the manufacturer’s guidelines with small modifications (Bionano Prep SP Frozen Human Blood DNA Isolation Protocol, Bionano Genomics #30246). In order to preserve DNA integrity and prevent clotting of the samples, DNA stabilizer was added to heparinized samples before or after freezing and some samples were additionally filtered with 100μm cell strainer (pluriStrainer Mini 100μm, pluriSelect) by centrifugation for 5 minutes at 400 x g. White blood cells (WBCs) were counted with HemoCue (Radiometer Benelux) and 1.5M cells were used for the DNA isolation protocol. Cells were pelleted (2,200 x g, 2min) and after removing the supernatant the cell pellet was resuspended in Proteinase K and RNAse (for bone marrow aspirates only). Following this, to release the gDNA, LBB lysis-buffer was added and the samples were mixed using HulaMixer (ThermoFisher Scientific). After PMSF treatment (Sigma-Aldrich), Nanobind disks were placed on each sample solution and isopropanol was added. Samples were then mixed using HulaMixer to bind the released gDNA onto the disks. After washing, the disks were transferred to fresh tubes and the gDNA was eluted from the disks. Finally, the gDNA was mixed and equilibrated overnight at room temperature to facilitate DNA homogeneity.

DNA quantification was carried out using Qubit dsDNA assay BR kit with a Qubit 3.0 Fluorometer (ThermoFisher Scientific). For 47/48 (97.9%) of the samples, the concentrations were above or equal 35 ng/μL as recommended and for 46/48 (95.8%) of the samples the coefficient of variation (CV) was <0.3 (three measurement points from each DNA sample), fulfilling the recommended criteria to perform the following gDNA labelling reaction (Supplementary Table 1).

### Labeling of UHMW gDNA and chip loading

The UHMW gDNA labelling was performed following the manufacturer’s guidelines using the Bionano Prep Direct Label and Stain (DLS) Protocol. Briefly, 750ng purified UHMW DNA was labelled with DL-green fluorophores using the Direct Labeling Enzyme (DLE-1) chemistry, followed by Proteinase K digestion (Qiagen) and DL-green cleanup using two membrane adsorption steps on a microplate. Finally, the labelled samples were homogenized by mixing with HulaMixer and stained overnight (Bionano DNA stain reagent) at room temperature, protected from light, to visualize the DNA backbone.

DNA quantification was carried out using Qubit dsDNA assay HS kit with a Qubit 3.0 Fluorometer (ThermoFisher Scientific). 40/48 (83.3%) of the labelled samples had concentrations within the recommended values of 4-12 ng/μl for both measurement points, 7/48 samples had one measurement >12 ng/μl, and only one sample had both measurements below 4 ng/μl (Supplementary Table 1).

### Data collection

Labelled gDNA samples were loaded on 3x 1300 Gb Saphyr chips (G2.3) and imaged by the Saphyr instrument. Each flowcell was run on maximum capacity to generate 1300 Gb of data per sample using Hg19 as the reference for real time quality control assessment.

### Assemblies and variant calling

*De novo* genome assemblies, variant calling and Rare Variant Pipeline were all performed via Bionano Access software (v1.4.3) using the Bionano Tools version 1.4.3 for assembly and variant calling and RefAligner v10020 for the Rare Variant Pipeline (RVP). For the current manuscript, only data from the RVP was used. For data filtering, a customized filter was created using the following confidence scores: Insertion: 0, Deletion: 0, Inversion: 0.01, Duplication: -1, Translocation: 0.01 (low stringency: filter set to 0), Copy Number: 0.99 (low stringency: filter set to 0). Per sample, prefiltered data were downloaded as .csv files for SVs and CNVs separately. These .csv files were used to determine the numbers and types of aberrations per sample (Table 1, Supplementary Table 2). ‘Whole genome CNV’ views were only enabled in the latest Bionano Access software version 1.5.

**Table 1:** Average numbers of SVs and CNVs with the recommended confidence filters.

| **<u>SV calls using the high confidence filter settings</u>** |  |  |  |  |
| --- | --- | --- | --- | --- |
|  | **Sum in all samples** |  | **Average per sample** |  |
|  | *all SVs* | *rare SVs* | *all SVs* | *rare SVs* |
| *Total* | 71247 | 2178 | 1454.0 | 44.4 |
| *Insertion* | 31452 | 711 | 641.9 | 14.5 |
| *Deletion* | 30981 | 989 | 632.3 | 20.2 |
| *Inversion* | 3633 | 70 | 74.1 | 1.4 |
| *Duplication* | 4701 | 246 | 95.9 | 5.0 |
| *Translocation interchromosomal* | 349 | 95 | 7.1 | 1.9 |
| *Translocation intrachromosomal* | 131 | 67 | 2.7 | 1.4 |
| **<u>Rare SVs overlapping genes (within 12kb window)</u>** |  |  |  |  |
|  | **Sum in all samples** |  | **Average per sample** |  |
| *Total* | 1254 |  | 25.8 |  |
| *Insertion* | 345 |  | 7.0 |  |
| *Deletion* | 557 |  | 11.4 |  |
| *Inversion* | 45 |  | 0.9 |  |
| *Duplication* | 192 |  | 3.9 |  |
| *Translocation interchromosomal* | 67 |  | 1.5 |  |
| *Translocation intrachromosomal* | 48 |  | 1.0 |  |
| **<u>CNV calls (non-masked only)</u>** |  |  |  |  |
|  | **Sum in all samples** |  | **Average per sample** |  |
| *Gain (called duplication in the file)* | 451 |  | 9.0 |  |
| *Loss (called deletion in the file)* | 2993 |  | 53.4 |  |
| *Putative amplifications (CN>3)* | 13 |  | 0.3 |  |

### Data comparison

For comparison of the Bionano data to the standard of care workflow, each pre-filtered csv file was investigated for the presence of the known aberrations (from karyotyping, CNV-microarray, and/or FISH) (Table 2). Only clinically relevant, reported SVs were taken into account. SVs with a variant allele fraction (VAF) of <10% (equivalent to the presence of SVs in 20% of cell fraction, as determined by CNV-microarray CNV profile or aberrant metaphase count by karyotyping) were excluded for the purpose of this study. Potential newly identified SVs are mentioned for occasional cases but the investigation of those is beyond the scope of this manuscript.

**Table 2a:**
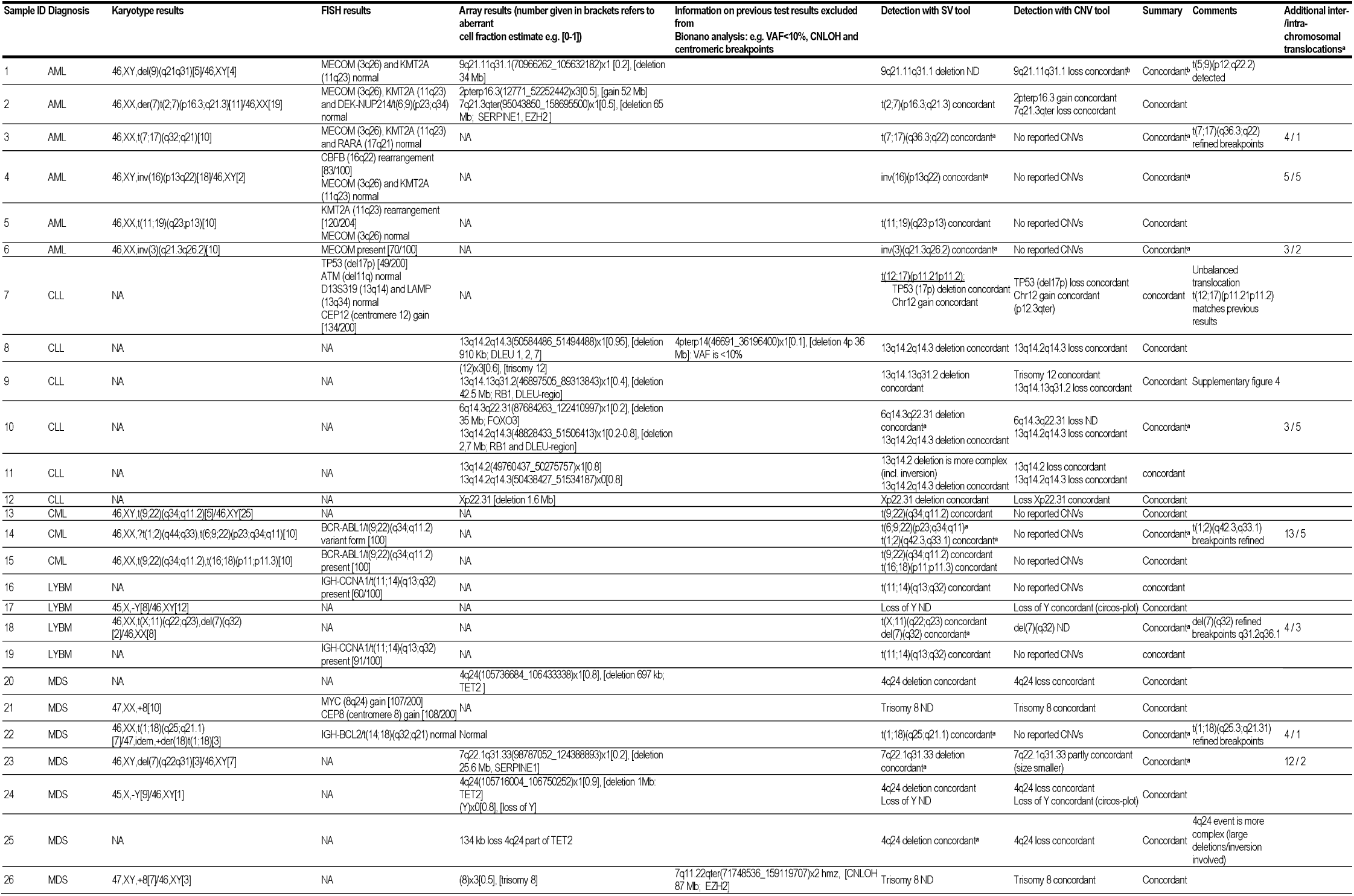
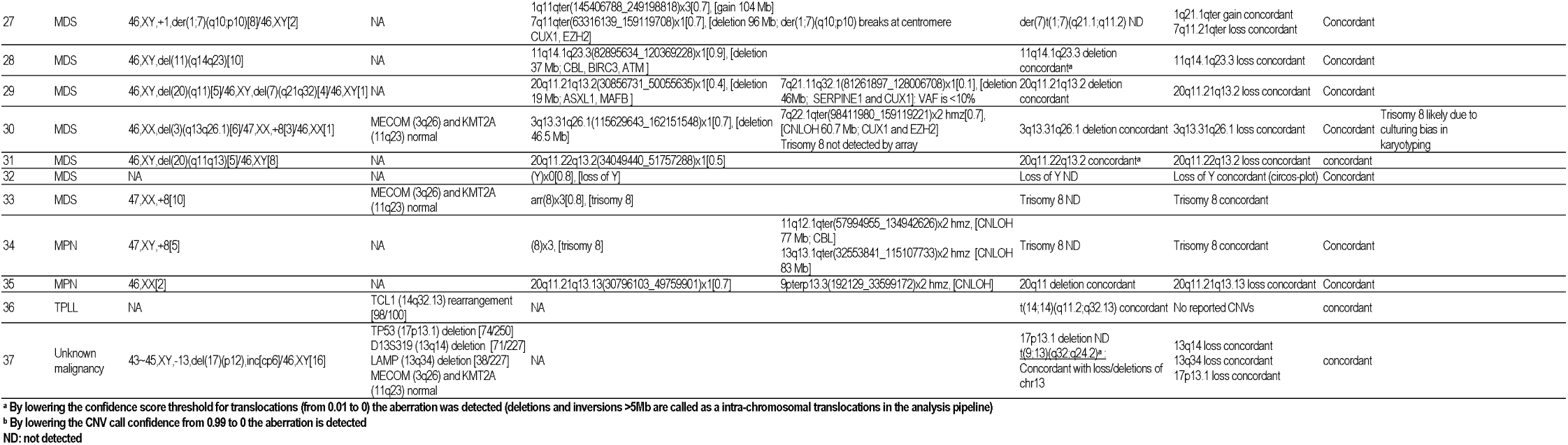
Comparison of previous findings (karyotyping, CNV-microarray and FISH) with genome imaging results for simple cases

**Table 2b:**
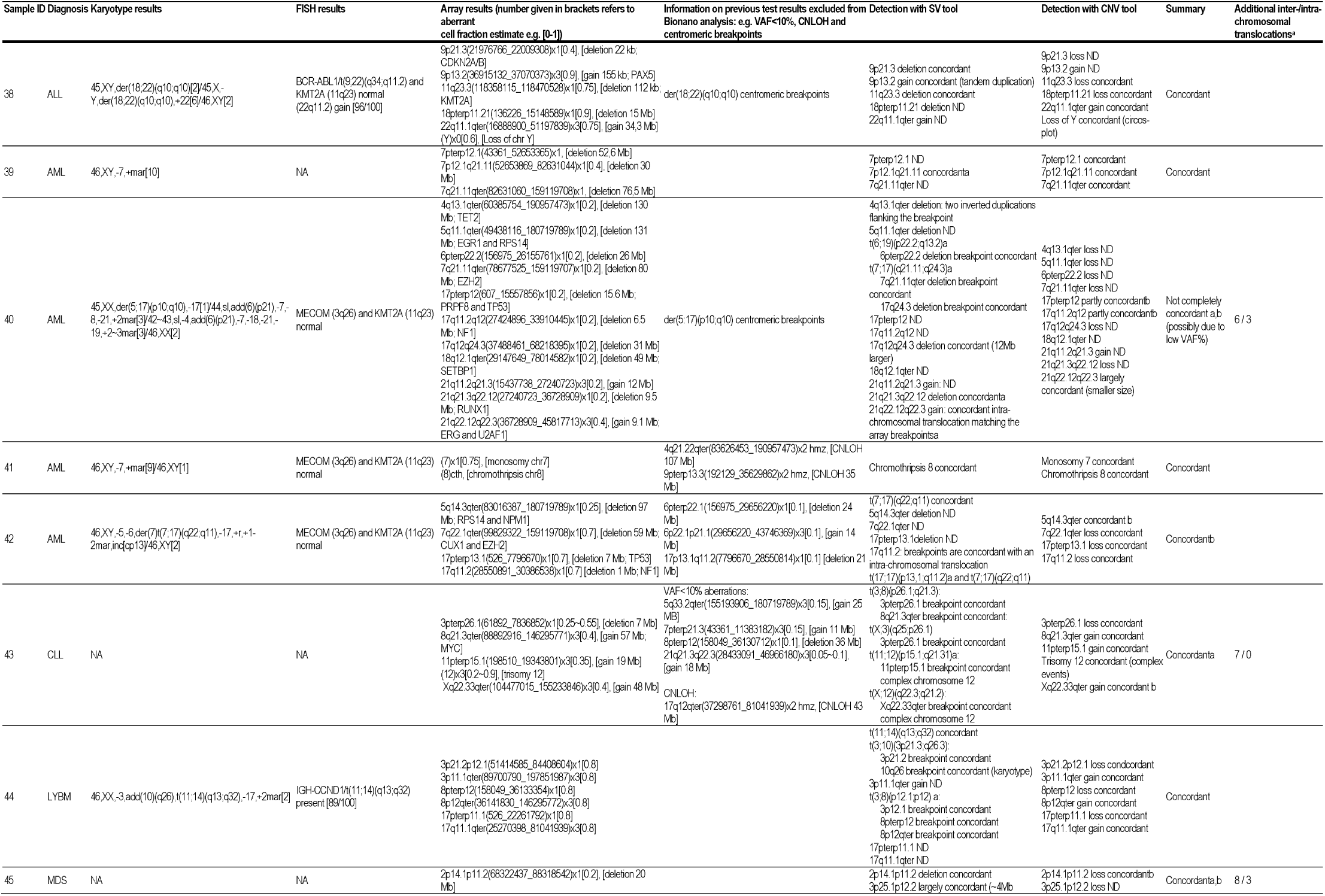
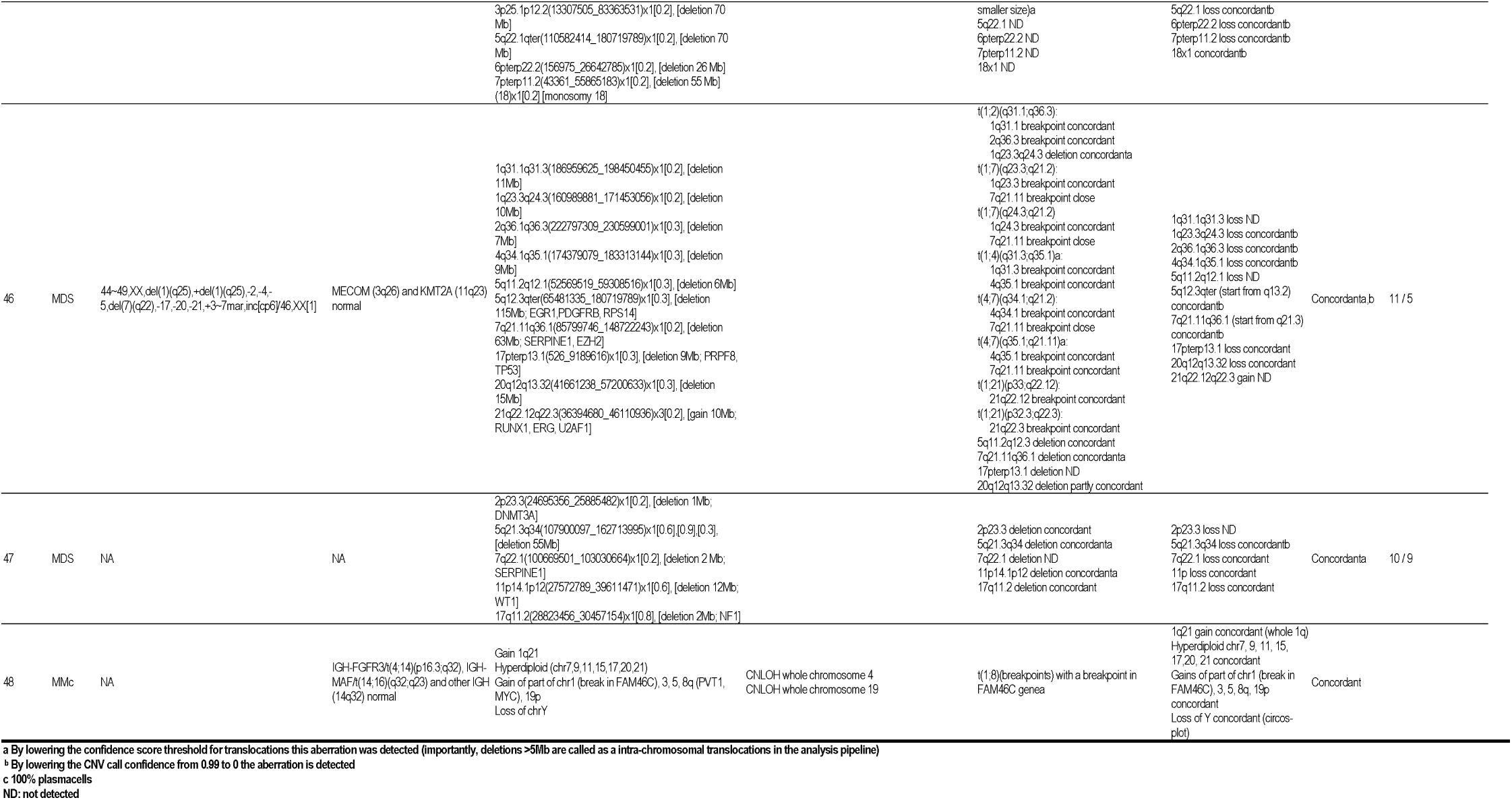
Comparison of previous findings (karyotyping, CNV-microarray and FISH) with genome imaging results for complex cases

### Terminology

Terminology as used in this article is based on 2 different algorithms that are incorporated in the Rare Variant pipeline, one for SVs and one for CNVs. Consequently, terminology is slightly different than commonly used in cytogenetics laboratories.

The SV tool calls insertions, deletions, duplications, inversions, inter- and intrachromosomal translocations.^20^ Intrachromosomal translocation breakpoints involve regions with a minimum distance of 5Mb from each other on the same chromosome, meaning that also deletions or inversions > 5Mb are called as intrachromosomal translocations.^20^ CNVs are instead detected based on coverage depth information using a copy number analysis pipeline embedded in the Rare Variant Pipeline. The copy number tool identifies fractional copy number changes and chromosomal aneuploidy events. ^20^

For the purpose of this study, we consider concordance of genome imaging with previous findings (CNV-microarray) whenever the same event is detected, even though the size or breakpoints of an SV/CNV may differ slightly (Table 2).

### Karyotyping

Bone marrow samples were cultured for 24 and 48 hours, respectively, in RPMI1640 medium supplemented with 10% fetal calf serum and antibiotics. After hypotonic treatment with 0.075M KCl and fixation in methanol/acetic acid (3:1) microscopic slides (GTG-banding) were prepared. Chromosomes were G-banded using trypsin and Giemsa and at least 20 metaphases were analyzed in case of a normal karyotype, and at least 10 in case of an abnormal karyotype. Karyotypes were described according to the standardized ISCN 2016 nomenclature system.

### FISH analysis

Standard cytogenetic cell preparations were used for fluorescence in situ hybridization (FISH). FISH was performed using commercially available probes according to the manufacturer’s specifications (Abbott Molecular, Des Plaines, Illinois). At least 200 interphase nuclei were scored by two independent investigators.

### CNV-microarray

CNV-microarray analysis was carried using the CytoScan HD array platform (ThermoFisher Scientific, Waltham, USA). Hybridizations were performed according to the manufacturer’s protocols. The data were analyzed using the Chromosome Analysis Suite software package (ThermoFisher Scientific, Waltham, USA), using annotations of genome version GRCh37 (hg19). Aberrations were described according to ISCN 2016.

## Results

### Samples included

All leukemia samples (n=48) in this study were first analyzed with the standard of care workflow, followed by the analysis of residual material on the Bionano Saphyr system to detect diagnostically relevant (*i.e*. reported) structural variants/chromosomal aberrations. We chose a combination of myeloid and lymphoid neoplasms (AML, MDS, CML, CLL, ALL, MM, MPN, T-PLL, LYBM) with an abnormal cytogenetics report to represent a broad set of clinically relevant SVs and CNVs. These are representative for the most common referrals to our clinic, with an estimated yearly number of 1,800 samples. Based on the diagnostically reported aberrations, the 48 samples were classified into two different groups: 37 samples with <5 aberrations (categorized as simple cases) and 11 samples with ≥5 aberrations or an unspecified marker chromosome (categorized as complex) (Table 2).

### Genome imaging results and SV/CNV calling

The optical mapping of these 48 leukemia genomes resulted in an average of 280-fold effective coverage (+/- 51.17), with an average label density of 14.6/100kb (+/- 1.62), a map rate of 71% (+/- 9.33) and an average N50 (>150kb) of 263kb (+/- 0.03) (Supplementary Table 1). In total, we identified 71,247 SVs and 7,301 CNVs in 48 leukemia samples (Supplementary Tables 3 and Supplementary Tables 4). Per sample, on average 1,454 total SVs were detected, comprising 642 insertions, 632 deletions, 74 inversions, 96 duplications, 7 interchromosomal translocations and 3 intrachromosomal translocations. Filtering these SVs for rare variants only, where rare is defined as variants that are not present in a cohort of 57 control samples, resulted in 2178 SVs in total (Supplementary Table 5). Each sample showed 44 rare SVs on average, of which 15 were insertions, 20 deletions, 1 inversion, 5 duplications, 2 interchromosomal translocation and 1 intrachromosomal translocation (Table 1, for examples see Figure 1 and Figure 2).

**Figure 1:**
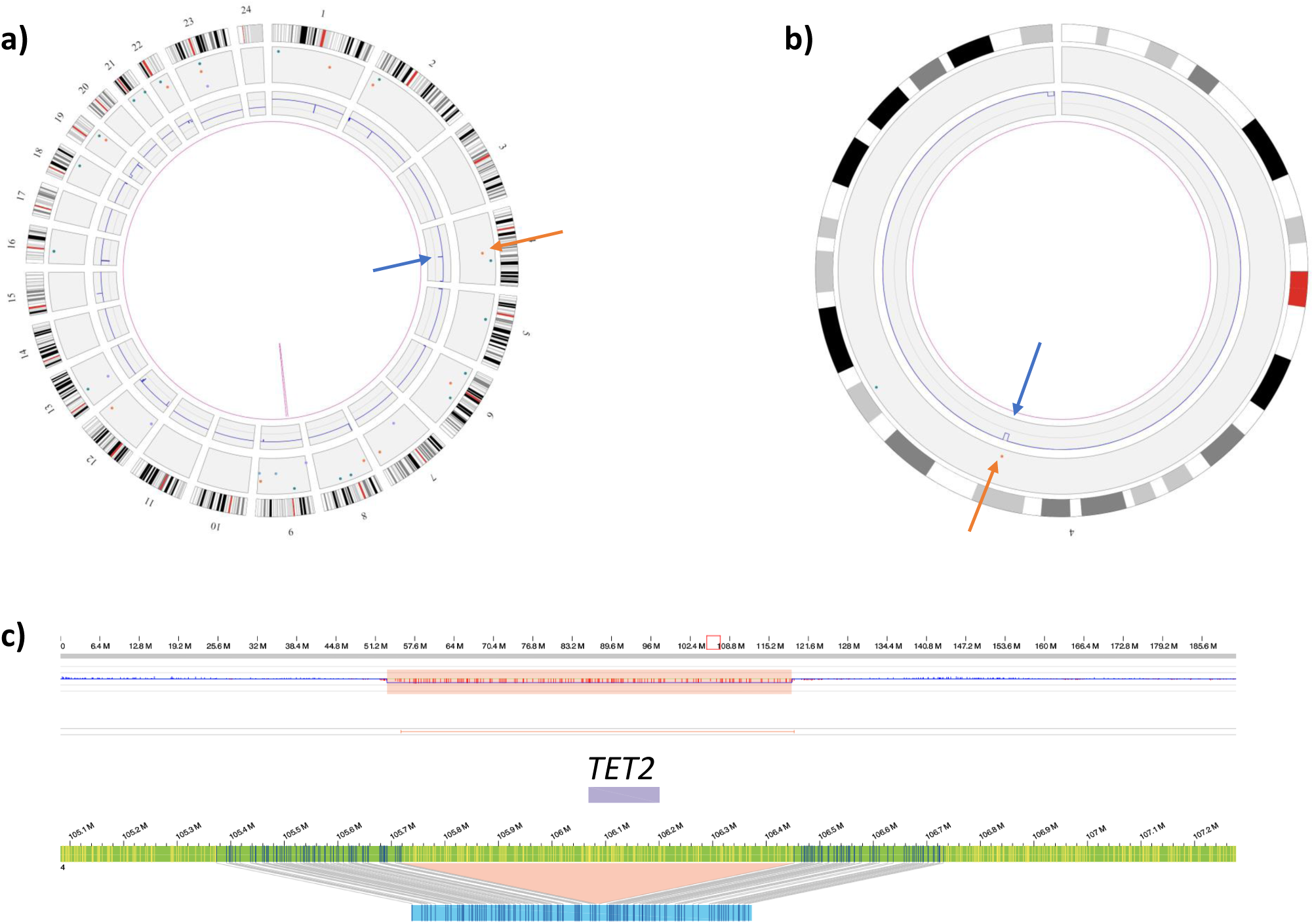
Example of Bionano output for one sample with a deletion (sample 20). Illustration of a known deletion spanning *TET2*, called by Bionano RVP. This deletion was previously called by CNV-microarray (boundaries: chr4:105,736,684-106,433,338, total size:697kb). This deletion is called by both the SV- and the CNV- tool. **a)** Whole genome circos plot showing a total of 47 rare SVs, of which 24 insertions, 12 deletions, 3 inversions, 7 duplications, and 1 intrachromosomal translocation. The *TET2* deletion is pointed out in the CNV track (blue arrow) and the SV track (orange arrow). **b)** Zoom-in circos plot of chromosome 4 only, showing the *TET2* deletion called by the SV and CNV tool. **c)** Navigation from the circos plot to the ‘chromosome maps view’ is enabled by selecting the respective chromosome or SV call, guiding to the map that supports the deletion. The lower track shows the SV deletion call from the map, specifying the breakpoint to chr4:105,717,890-106,452,233 (size: 727.7kb). The upper track illustrates the CNV call from an independent tool.

**Figure 2:**
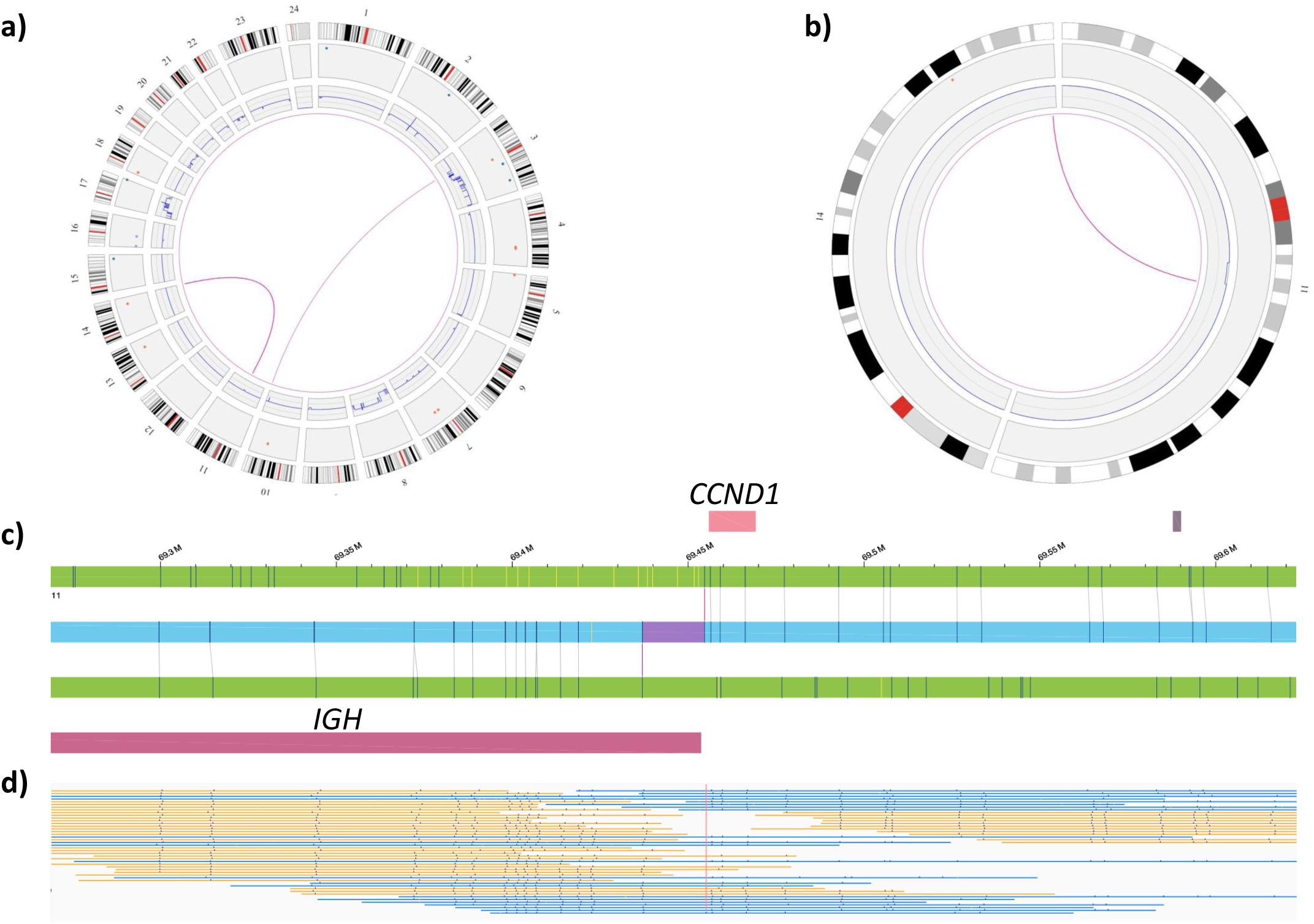
Example of Bionano output for one sample with a balanced translocation (sample 44). Illustration of a known translocation t(11;14) called by Bionano RVP. **a)** Whole genome circos plot showing 2 translocations (t(11;14) and t(3;10), purple lines). **b)** Zoom-in circos plot of ‘affected’ chromosomes only, enabling a more detailed visualization of t(11,14). **c)** Navigation from the circos plot to the ‘chromosome maps view’ is enabled by clicking the purple line, guiding to the map that supports the translocation. The selected example shows the translocation breakpoints t(11;14), mapping to genes *CCND1* (chr.11) and *IGH* (chr.14). **d)** Aberrant molecules supporting the translocation.

For CNVs, filtering for rare variants is not available yet. However, the RVP analysis automatically masks regions of the genome with unusually high variance in their relative coverage across control datasets (including centromeric and telomeric regions), assuming that these high variance regions may be regions of high CNV occurrence in normal healthy individuals.^21^ For the total of 7,301 CNV calls, 3,445 CNV calls (451 gains, 2,993 losses) were left after masking. Per sample, this lead to an average of 9 (range: 0-64) non-masked gains and 53 (range: 4-575) non-masked losses (Table 1). Segmentation of large CNVs, partial trisomies or monosomies inflates these numbers, similar to findings in CNV-microarrays.

### Improved filter settings lead to 100% true positive rate for known aberrations for simple cases

By applying our customized filtering settings (see Materials and Methods), all diagnostically reported aberrations were identified by a combination of SV and CNV outputs, with the exception of 10 events in 9/37 simple cases, resulting in the correct identification of 78% (36/46) of diagnostically reported aberrations with variant allele fraction >10% (excluding CN-LOH) (Table 2). Nine out of the 10 aberrations that escaped the initial filtering were detected by lowering the confidence threshold for translocations from 0.01 to 0, although not all of these aberrations were actually translocations (see Terminology in Material and Methods). Lowering the confidence filters for translocations resulted in the calling of on average 9 (range: 5-18) additional aberrations per sample (Table 2). Independent validations are needed to determine the true-positive of false-positive character of the additional translocations. The one missing aberration that could not be rescued by a lower translocation confidence filter concerned a sample with bad quality (N50: 205kb, map rate: 53%). Nevertheless, even this one could be unraveled by reducing the CNV confidence to 0, importantly resulting in a total of 100% detection rate of diagnostically reported findings.

Of note, aneuploidies of sex chromosomes were clearly visible in CNV profiles and circos plots, but not called due to the absence of a counting for sex-chromosomes in the current RVP analysis.

### Correlation of known clinically relevant findings with Genome Imaging in complex cases

Ten of the 11 complex cases showed full concordance with previous findings (Table 2). Of those ten, five required the lowered filter settings in order to identify all respective aberrations (two required lower settings for SVs, one for CNVs, and two for both) (Supplementary Figure 1 and Supplementary Figure 2). Only in one case we did not observe full concordance with previous findings. However, this represents a case with very complex aberrations of which we still identify the majority. All missed aberrations had a previously estimated VAF of ∼10% (20% estimated aberrant cell fraction by CNV-microarray). Next to still identifying the vast majority of known aberrations, we observed that genome imaging likely reveals the true underlying nature of aberrations. For several cases breakpoints of gains and losses identified by CNV-microarrays match the translocation breakpoints identified by genome imaging or refined previously known translocations from karyotyping.

Very interestingly, even a complex chromothripsis structure was resolved unambiguously (case 41, Table 1, Figure 3). In addition, for several of the complex cases it seems that the clinically detected rearrangements are even more complex than previously seen, e.g. additional translocations were identified or marker chromosomes of unknown origin were resolved (Table 2).

**Figure 3.**
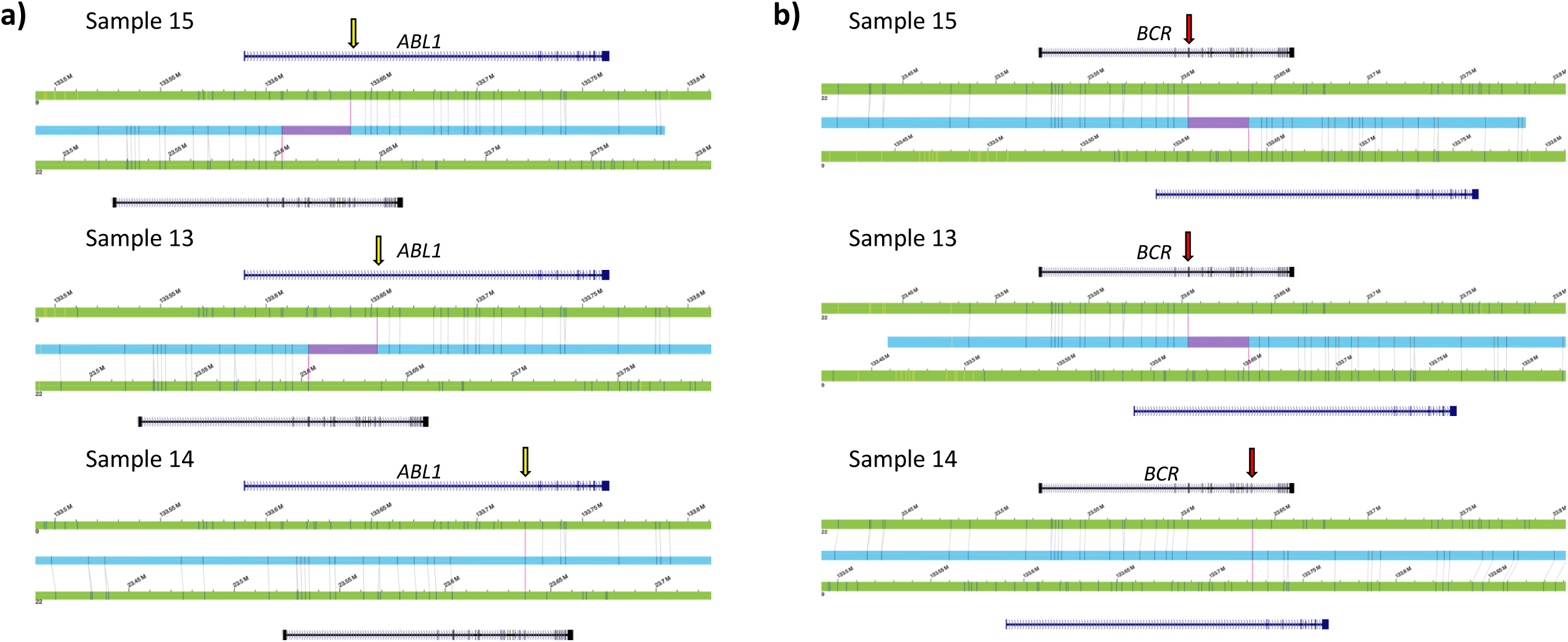
Resolving chromothripsis structures. A complex chromosome 8 chromothripsis structure was resolved unambiguously in an AML sample (sample 41). **a)** The circos plot illustrates the shattering of chromosome 8, called as intra-chromosomal translocations. In addition, a monosomy of chromosome 7 was identified. **b)** Zoom-in to chromosome 8, showing the aberrant CNV profile (top) and maps of the rare variant pipeline (bottom).

### Novel findings by genome imaging – translocations leading to potential fusions

Next to the very high concordance for diagnostically reported aberrations, genome imaging also finds novel aberrations. While this was not the major purpose of this study, we aimed to share all SVs and CNVs with the field (Supplementary Tables 3 and Supplementary Tables 4) that may enable future discoveries.

As genome imaging has a unique ability to identify balanced aberrations with high resolution, we focused our analysis on potential fusion genes that resulted from interchromosomal translocations, due to their known importance as (novel) leukemia driver mutations. From the list of rare SVs (n=2,178, Supplementary Table 5), 95 presented as rare interchromosomal translocations. Of those, 23 were unique calls leading to potential gene fusions (based on Bionano’s annotations) (Supplementary Table 6). Four of those were known, previously diagnostically reported fusions, including the only recurrent fusion which was reported in 3 independent samples affecting the *BCR-ABL1* fusion gene (Figure 4) and one known fusion of *KMT2A*-*ELL*. Of the remaining potential fusions no recurrent events were observed (Supplementary Table 6), and none was reported previously (COSMIC catalogue of somatic mutations in Cancer). Of potential interest are putative fusions of *RUNX1-AGBL4* and of *BCR-EXOC2* (Supplementary Figure 3), for both of which one of the two fusion partners is well known from other fusion genes in leukemia. The latter was another BCR-fusion from a case with a three-way translocation involving the Philadelphia chromosome and BCR-ABL1 fusion.

**Figure 4.**
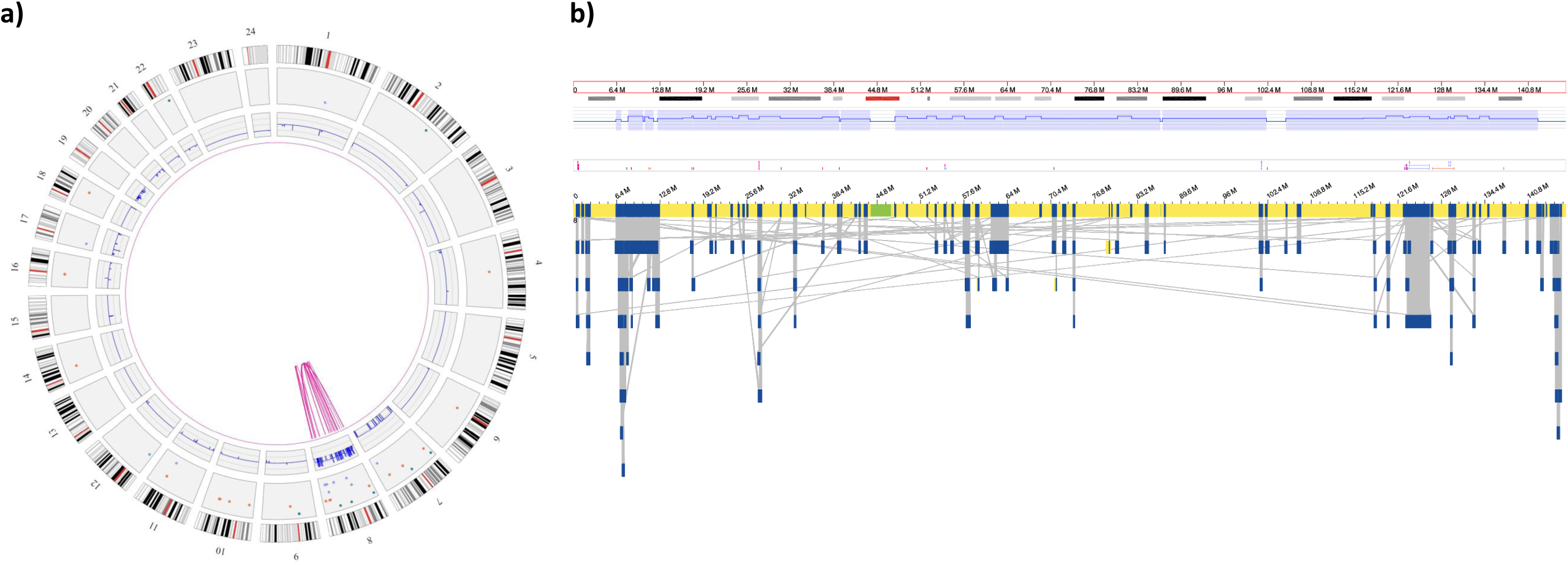
Recurrent BCR-ABL1 fusion genes. Data analysis identified 3 recurrent translocations, leading to *BRC-ABL1* fusion genes in 3 CML cases. The resolution of optical mapping allowed to identify possibly 2 different BCR breakpoints in the three samples (chromosome 22), however they differ by only one label. We were also able to distinguish three different breakpoints in *ABL1* (chromosome 9), all mapping to intron 1. Translocation breakpoints are indicated by yellow (*ABL1*) and red (*BCR*) arrows.

### Novel findings by genome imaging – other SVs

To explore the ability to detect smaller SVs that may escape traditional methods (e.g. CNV-microarrays), we checked all rare deletions and insertions calls <10kb in size (n=963, Supplementary Table 5). Of those, 26 calls overlap with well-established cancer genes (Supplementary Table 7).^22^ Three genes show recurrent small insertions and deletions. Most interesting candidates are a 6kb deletion and a 3.3 kb insertion affecting *NOTCH1* in sample 14 (CML) and 30 (MDS) respectively. Additional validations for these novel findings that usually also escape calling from CNV-microarrays are warranted.

## Discussion

Genome imaging on the Bionano Genomics’ Saphyr system (Bionano Genomics, San Diego) relies on a high-throughput comparison of distance and pattern of fluorescent labels on long DNA molecules >150 kb to the respective distance and patterns in a given references sequence (e.g. hg19 or hg38 after *in silico* digestion). Only recently, increased throughput, lowered costs, and improved resolution allow for the usage of this technology for structural variant detection in clinical relevant human applications. The identification of structural variants is key for the diagnostics of genetic disorders. Recent work from Barseghyan et al.^10^ illustrates this by showing how genome imaging correctly diagnoses Duchenne Muscular Dystrophy from clinical samples. In a another study a prostate tumor sample was profiled by comparing the cancer sample with matched blood by genome imaging.^23^

In the current study, we aimed to investigate whether the optical mapping technology would be suited to replace karyotyping, CNV-microarray and FISH as single diagnostic test for hematological diseases. Therefore, we compared previously reported diagnostic data from 48 leukemia samples with data generated by genome imaging. All samples performed according to specifications, with a label density of 14.6/100kb and an N50 molecule length (>150kb) of 263kb resulting in 280x genome coverage.

### Genome imaging for simple cases

Most remarkably, we were able to identify all previously reported aberrations with a VAF>10% (SVs and CNVs) in the simple cases. Identification of 100% of the aberrations however required to lower the stringency settings used for filtering, mainly for translocations, and lead to the additional calling of on average 9 aberrations per sample. Whether these additional aberrations present true or false positives still needs to be determined, and a combination of both is likely. Manual inspection of several of those translocations showed that they often were supported by many high quality molecules, suggesting that they might be real and that readjustment of the confidence values is needed for future software upgrades. Recurrent intrachromosomal translocations on chromosome 9, that were identified in some cases with lowered translocation thresholds, instead lead to the hypothesis that other additional findings may be false positives – or represent the true complexity of that locus and may indicate issues in the human reference genome.

### Genome imaging for complex cases

For the complex cases we also observe a very high concordance with previous findings. Only in one case we did not observe full concordance. However, the previously estimated VAF of ∼10% (20% estimated aberrant cell fraction by CNV-microarray) may not have been accurate, assuming that an actual lower aberrant cell fraction is not unlikely. If this was the case, this would be beyond the scope of our study. Surprisingly we still detect the vast majority of all aberrations despite the low VAFs.

Especially for complex cases, a key benefit of genome imaging became apparent: Only a combined assay that enables the detection of (almost) all aberrations in one test is able to unravel the true underlying architecture of complex genomic re-arrangements. Now we observe that several of the gains and losses identified by CNV-microarrays match the translocation breakpoints identified by genome imaging. Previously this always required karyotyping as translocations cannot be identified with CNV-microarrays. But even karyotyping may miss events or may not identify exact breakpoints, as also observed for several cases of our study (Table 2). Occasionally, karyotyping is even impossible, e.g. when no metaphase chromosomes were obtainable.

### Novel findings by genome imaging

Next to the identification of previously known aberrations, we also checked for novel ones although this was not the main purpose of this study. We especially concentrated on translocations leading to potential novel fusion genes, since those are important drivers for cancer development and discovery of new drivers can lead to important biological insight and potential new treatment possibilities.^24^ Traditionally, fusion-gene mapping used methods like SKY-FISH followed by FISH and PCR to identify one or both fusion partners of recurrent translocations.^25^ Nowadays, short-read genome sequencing can successfully identify different kind of variations, including translocations leading to novel fusion genes.^24; 26; 27^ Emerging long-read genome sequencing technologies make identification of SVs even easier,^28-31^ and show potential to unravel complex events like chromothripsis.^30; 32^ There have also been successful applications of RNAseq for the detection of fusion genes in leukemia,^27; 33^ which has however not entered the clinical setting yet and the sensitivity for low VAF needs to be shown. Unfortunately, many of these new technologies require a combination of optimized research analysis tools and a dedicated team of skilled bioinformaticians, and therefore are often applied by large consortia or large-scale genome sequencing centers only. Despite the advantages and potential advantages of genome sequencing, sequencing technologies remain limited by interspersed repeat elements which are longer than the sequence reads and therefore do not allow unique mapping, which masks most SVs in short read sequencing and some in long read sequencing.

Genome imaging now offers the possibility for a rather easy and direct identification of such fusions, as well as an easy detection of inversions that were likewise difficult to identify until now. Its independence of sequence context in combination with the ultra-long molecules enables the analysis of even the most complex regions of the genome.

Another type of novel findings are small SVs that usually escape detection by classical means. To get a first expression, we filtered for aberrations smaller than 10kb, and for example identified two different SVs affecting *NOTCH1*, being a 6kb deletion and a 3.3 kb insertion in sample 14 (CML) and 30 (MDS) respectively. This is a potentially interesting finding, as NOTCH 1 is an important leukemia driver gene in CLL,^34^ but also emerges for other leukemia types.^35^ Whether these two SVs add anything meaningful to our two cases requires further investigations though.

### Resolution of genome imaging

We observe that for a wide range of structural variants genome imaging offers higher resolution compared to the standard technologies. Current resolutions and reporting criteria of standard-of-care technologies are 5Mb for karyotyping, and for CNV-microarrays 100kb-5Mb for leukemia specific regions and >5Mb for non-leukemia specific regions. For CNV-microarrays also copy-number neutral LOH (CNLOH) is reported when >10Mb and extending towards the telomere, just as mosaicism down to 10%. FISH can only detect specific rearrangements and fusion genes. Genome imaging in combination with the Rare Variant Pipeline instead allows the detection of insertions within a size-range of 5-50kb, deletions >5kb, translocations (or transpositions) >70kb, inversions >100kb and duplications >150kb.^20^ This higher resolution of genome imaging for all types of variants will allow the detection of smaller cancer-associated events, potentially leading to new insights and maybe even better treatment options.

### Current challenges and opportunities

Although genome imaging comes with a lot of advantages, there are also limitations to this new technology. First, the detection of Robertsonian translocations, or any other balanced translocations with breaks in the repeat regions surrounding the centromere, is not possible yet due to missing labels for the centromeres. Second, we did not yet include the detection of events with a VAF <10% systematically. Another study, including systematic dilution series, would be required to test detection limits of lowest level somatic aberrations. Third, CN-LOH identification is not enabled yet. We have however analyzed two exemplary samples with previously identified large LOH regions (sample 26 and 41, 87 and 107 Mb respectively), and within the homozygous regions 88% (37/42) and 89% (49/55) of SV calls were called as homozygous by the *de novo* assembly algorithm (RVP does not support genotyping of called events). We believe that with some improvements this can allow to call LOH at least for larger regions spanning several Mb. We believe that calling missing or additional labels, due to a (common) SNP in the 6mer recognition motif, will allow improved ‘genotyping’ and could further improve LOH calling.

Concerning the detection of events with a VAF <10%, which were too rare in our cohort to draw major conclusions, the RVP tool only requires a default minimum of 3 molecules showing the identical SV^20^, hence lower VAFs shall be identified when higher coverage is enabled. We also anticipate that higher throughput, and usage of the 2^nd^ laser that is already included in the Saphyr instrument but not actively used yet, will enable the use of a 2^nd^ dye that would allow for sample-barcoding and pooling on the same flowcell in future. This should enable at least to pool 2 samples that have been differentially labeled, but the average of ∼50 labels per molecule should also allow ratio labelling with two dyes, allowing to pool three or more samples, as was shown previously for FISH.^36^

Turnaround time is critical for clinical implementation of this test, and the current amount of data especially for high coverage somatic SV detection makes this challenging. We already observed a dramatic improvement in analysis time for RVP vs. *de novo* assembly, but further improvements are necessary. One might need to think about applying the RVP tool only to a defined set of genes or regions of interest, as was recently demonstrated for FSHD,^37^ which would dramatically speed up analysis and reduce number of aberrations that need to be considered. Furthermore, we would prefer one combined report for the SV and CNV outputs coming from two different algorithms, and think that data would be best visualized by a composite ideogram style, such as described in ^38^, and by ‘genome-wide CNV profiles’ similar to the CNV-microarrays that are now available within the most recent update of the Bionano software (Supplementary Figure 4). Finally, other very intriguing new developments include the potential for ‘phased’ detection of SVs and methylation,^39^ which can offer additional value for cancer research and diagnostics.^40^

While we can show a true positive rate of 100% for simple cases, our study is not yet suited to determine sensitivity and specificity. We identified additional aberrations in many cases, which either were not identified or not reported by standard of care tests. Full assessment of false-positive rates of genome imaging however would require orthogonal validations, some of which would not be trivial. But the overall numbers identified here led us to conclude that false-positive rates are very likely (very) low. This conclusion is also reflected by the analysis of 11 complex cases while we see strong concordance with previous results, usually genome imaging allows better resolution and a more complete picture of complex aberrations.

In summary, the data presented in this manuscript and the promising future improvements of the technology convince us that the use of high resolution genome imaging for diagnostic purposes will be feasible in near future. Thereby, optical mapping has the potential to replace existing cytogenetics analyses and may become the one generic test for all (molecular) cytogenetic applications, thereby being highly complementary to existing sequencing based technologies.

## Supporting information

Supplemental Table 1: Technical performance of genome imaging

Supplemental Table 2: Absolute numbers of all SVs and CNVs with the high confidence filters per sample

Supplemental Table 3: All 71247 SVs called by high confidence filters

Supplemental Table 4a: All CNVs in all samples, Supplemental Table 4b: All 3445 non-masked CNVs

Supplemental Table 5: All rare 2178 SVs in all called by high confidence filters

Supplemental Table 6: Possible fusion-genes resulting from high-confidence inter-chromosomal translocations

Supplemental Table 7: All rare SVs (<10kb) overlapping with known COSMIC cancer genes

## Acknowledgements

We are thankful to the Department of Human Genetics (especially Helger Ijntema, Lisenka Vissers, Marjolijn Ligtenberg, Marcel Nelen, Han Brunner) for providing support and critical feedback. We are grateful to the Radboud UMC Genome Technology Center for infrastructural and computational support. We would like to acknowledge support from scientists and staff at Bionano Genomics including Alex Hastie, Andy Pang, Lucia Muraro, Kees-Jan Francoijs, Sven Bocklandt, Yannick Delpu, Mark Oldakowski, Ernest Lam, Thomas Anantharaman, Scott Way, Henry Sadowski, Amy Files, Carly Proskow. Financial support (to AH) was given by the European Union’s Horizon 2020 research and innovation program Solve-RD (grant agreement No 779257) and the Dutch X-omics initiative, which is partly funded by NWO, project 184.034.019. TM was supported by the Sigrid Jusélius Foundation.

## Web Resources

Bionano Access™: https://bionanogenomics.com/support/software-downloads/#bionanoaccess Cosmic Catalogue of Somatic Mutations in Cancer: https://cancer.sanger.ac.uk/cosmic

## Abbreviations

ALL: acute lymphoblastic leukemia
AML: acute myeloid leukemia
BMA: bone marrow aspirates
CLL: chronic lymphocytic leukemia
CML: chronic myeloid leukemia
CNLOH: copy number neutral loss-of heterozygosity
CNV: copy number variant
CV: coefficient of variation
DLE-1: direct Labeling Enzyme-1
DLS: direct Label and Stain
EDTA: ethylenediaminetetraacetic acid
FISH: fluorescence *in situ* hybridization
FSHD: facioscapulohumeral dystrophy
gDNA: genomic DNA
i.e.: id est (that is)
LYBM: lymphoma in bone marrow
MDS: myelodysplastic syndrome
MM: multiple myeloma
MPN: myeloproliferative neoplasm
NGS: next generation sequencing
NIPT: non-invasive prenatal testing
PCR: polymerase chain reaction
PMSF: phenylmethylsulphonyl fluoride
RVP: rare variant pipeline
SD: standard deviation
SNP: single nucleotide polymorphism
ssDNA: single stranded DNA
SV: structural variant
TPLL: T-cell prolymphocytic leukemia
UHMW: ultra-long high molecular weight
VAF: variant allele frequency
WBCs: white blood cells

## Supplemental Data description

Supplemental Data include 7 tables and 4 figures.

**Supplemental Figure 1. Examples of filter adjustment for CNV calling**. Similarly to SVs, lowering the stringency threshold for CNV calling to 0 resulted in the calling of previously not called CNVs. For example **a)** 5q22.1qter loss (VAF 10%) in sample 45 (complex, MDS): lower stringency threshold shows larger CNV region than higher stringency threshold (red bars). **b)** Monosomy 18 (VAF 10%) in sample 45 (complex, MDS): whole chromosome 18 loss only seen with lower stringency threshold (red bars). **c)** Loss of 5q21.3q34 (sample 47, MDS): two different VAFs% (30%,45%) detected by high confidence threshold, VAF of 15% only detected with lower stringency threshold.

**Supplemental Figure 2. Circos plots of all complex cases**. Left row shows circos plots with high stringency filters for SVs and CNVs, middle row shows circos plots with low stringency filters for SVs (translocations only) and right row shows circos plots with low stringency filters for CNV calling.

**Supplemental Figure 3. Fusion gene detection. a)** Putative fusions of *RUNX1-AGBL4* (sample 46) and b) putative fusion of *BCR-EXOC2* (sample 14), for both of which one of the two fusion partners is well known from other fusion genes in leukemia. The latter was identified in a CML sample with a three-way translocation involving the Philadelphia chromosome t(9;22;6).

**Supplemental Figure 4: Comparison of CNV-microarray data and genome imaging**. Comparison of CNV-microarray results with the new ‘whole genome CNV’ visualization that is enabled in the latest Bionano Access software v1.5, for two cases. **a)** Unbalanced translocation t(2;7)(p16.3;q21.3) in sample 2, shown with array (upper panel) and genome imaging data (lower panel), **b)** *Trisomy 12 and del13q14.13q31.2 in sample 9*, shown with array (upper panel) and genome imaging data (lower panel).

**Supplemental Table 1: Technical performance of genome imaging**

**Supplemental Table 2: Absolute numbers of all SVs and CNVs with the high confidence filters per sample**

**Supplemental Table 3: All 71247 SVs called by high confidence filters Supplemental**

**Table 4a: All CNVs in all samples**

**Supplemental Table 4b: All 3445 non-masked CNVs**

**Supplemental Table 5: All rare 2178 SVs in all called by high confidence filters**

**Supplemental Table 6: Possible fusion-genes resulting from high-confidence inter-chromosomal translocations**

**Supplemental Table 7: All rare SVs (<10kb) overlapping with known COSMIC cancer genes**

## References

1. Vissers, L., van Nimwegen, K.J.M., Schieving, J.H., Kamsteeg, E.J., Kleefstra, T., Yntema, H.G., Pfundt, R., van der Wilt, G.J., Krabbenborg, L., Brunner, H.G., et al. (2017). A clinical utility study of exome sequencing versus conventional genetic testing in pediatric neurology. Genet Med 19, 1055–1063.

2. Lo, Y.M., Corbetta, N., Chamberlain, P.F., Rai, V., Sargent, I.L., Redman, C.W., and Wainscoat, J.S. (1997). Presence of fetal DNA in maternal plasma and serum. Lancet 350, 485–487.

3. Lo, Y.M., and Lam, W.K. (2016). Tracing the tissue of origin of plasma DNA-feasibility and implications. Ann N Y Acad Sci 1376, 14–17.

4. Rack, K.A., van den Berg, E., Haferlach, C., Beverloo, H.B., Costa, D., Espinet, B., Foot, N., Jeffries, S., Martin, K., O’Connor, S., et al. (2019). European recommendations and quality assurance for cytogenomic analysis of haematological neoplasms. Leukemia 33, 1851–1867.

5. Mikhail, F.M., Heerema, N.A., Rao, K.W., Burnside, R.D., Cherry, A.M., and Cooley, L.D. (2016). Section E6.1-6.4 of the ACMG technical standards and guidelines: chromosome studies of neoplastic blood and bone marrow-acquired chromosomal abnormalities. Genet Med 18, 635–642.

6. Bocklandt, S., Hastie, A., and Cao, H. (2019). Bionano Genome Mapping: High-Throughput, Ultra-Long Molecule Genome Analysis System for Precision Genome Assembly and Haploid-Resolved Structural Variation Discovery. Adv Exp Med Biol 1129, 97–118.

7. Chan, S., Lam, E., Saghbini, M., Bocklandt, S., Hastie, A., Cao, H., Holmlin, E., and Borodkin, M. (2018). Structural Variation Detection and Analysis Using Bionano Optical Mapping. Methods in molecular biology 1833, 193–203.

8. Lam, E.T., Hastie, A., Lin, C., Ehrlich, D., Das, S.K., Austin, M.D., Deshpande, P., Cao, H., Nagarajan, N., Xiao, M., et al. (2012). Genome mapping on nanochannel arrays for structural variation analysis and sequence assembly. Nature biotechnology 30, 771–776.

9. Schwartz, D.C., Li, X., Hernandez, L.I., Ramnarain, S.P., Huff, E.J., and Wang, Y.K. (1993). Ordered restriction maps of Saccharomyces cerevisiae chromosomes constructed by optical mapping. Science 262, 110–114.

10. Barseghyan, H., Tang, W., Wang, R.T., Almalvez, M., Segura, E., Bramble, M.S., Lipson, A., Douine, E.D., Lee, H., Delot, E.C., et al. (2017). Next-generation mapping: a novel approach for detection of pathogenic structural variants with a potential utility in clinical diagnosis. Genome Med 9, 90.

11. Seo, J.S., Rhie, A., Kim, J., Lee, S., Sohn, M.H., Kim, C.U., Hastie, A., Cao, H., Yun, J.Y., Kim, J., et al. (2016). De novo assembly and phasing of a Korean human genome. Nature 538, 243–247.

12. Shi, L., Guo, Y., Dong, C., Huddleston, J., Yang, H., Han, X., Fu, A., Li, Q., Li, N., Gong, S., et al. (2016). Long-read sequencing and de novo assembly of a Chinese genome. Nat Commun 7, 12065.

13. Sarsani, V.K., Raghupathy, N., Fiddes, I.T., Armstrong, J., Thibaud-Nissen, F., Zinder, O., Bolisetty, M., Howe, K., Hinerfeld, D., Ruan, X., et al. (2019). The Genome of C57BL/6J “Eve”, the Mother of the Laboratory Mouse Genome Reference Strain. G3 (Bethesda) 9, 1795–1805.

14. Bickhart, D.M., Rosen, B.D., Koren, S., Sayre, B.L., Hastie, A.R., Chan, S., Lee, J., Lam, E.T., Liachko, I., Sullivan, S.T., et al. (2017). Single-molecule sequencing and chromatin conformation capture enable de novo reference assembly of the domestic goat genome. Nat Genet 49, 643–650.

15. Jiao, Y., Peluso, P., Shi, J., Liang, T., Stitzer, M.C., Wang, B., Campbell, M.S., Stein, J.C., Wei, X., Chin, C.S., et al. (2017). Improved maize reference genome with single-molecule technologies. Nature 546, 524–527.

16. Zook, J.M., Hansen, N.F., Olson, N.D., Chapman, L.M., Mullikin, J.C., Xiao, C., Sherry, S., Koren, S., Phillippy, A.M., Boutros, P.C., et al. (2019). A robust benchmark for germline structural variant detection. bioRxiv, 664623.

17. Mak, A.C., Lai, Y.Y., Lam, E.T., Kwok, T.P., Leung, A.K., Poon, A., Mostovoy, Y., Hastie, A.R., Stedman, W., Anantharaman, T., et al. (2016). Genome-Wide Structural Variation Detection by Genome Mapping on Nanochannel Arrays. Genetics 202, 351–362.

18. Chaisson, M.J.P., Sanders, A.D., Zhao, X., Malhotra, A., Porubsky, D., Rausch, T., Gardner, E.J., Rodriguez, O.L., Guo, L., Collins, R.L., et al. (2019). Multi-platform discovery of haplotype-resolved structural variation in human genomes. Nat Commun 10, 1784.

19. Hastie, A.R., Lam, E.T., Chun Pang, A.W., Zhang, X., Andrews, W., Lee, J., Liang, T.Y., Wang, J., Zhou, X., Zhu, Z., et al. (2017). Rapid Automated Large Structural Variation Detection in a Diploid Genome by NanoChannel Based Next-Generation Mapping. bioRxiv, 102764.

20. Bionano Genomics. (2019). Bionano Solve Theory of Operation: Structural Variant Calling. Document Number: 30110, Revision F.

21. Bionano Genomics. (2019). Introduction to Copy Number Analysis. Document Number : 30210, Document Revision: D.

22. Tate, J.G., Bamford, S., Jubb, H.C., Sondka, Z., Beare, D.M., Bindal, N., Boutselakis, H., Cole, C.G., Creatore, C., Dawson, E., et al. (2019). COSMIC: the Catalogue Of Somatic Mutations In Cancer. Nucleic Acids Res 47, D941–D947.

23. Jaratlerdsiri, W., Chan, E.K.F., Petersen, D.C., Yang, C., Croucher, P.I., Bornman, M.S.R., Sheth, P., and Hayes, V.M. (2017). Next generation mapping reveals novel large genomic rearrangements in prostate cancer. Oncotarget 8, 23588–23602.

24. Gao, Q., Liang, W.W., Foltz, S.M., Mutharasu, G., Jayasinghe, R.G., Cao, S., Liao, W.W., Reynolds, S.M., Wyczalkowski, M.A., Yao, L., et al. (2018). Driver Fusions and Their Implications in the Development and Treatment of Human Cancers. Cell Rep 23, 227–238 e223.

25. Brockschmidt, A., Trost, D., Peterziel, H., Zimmermann, K., Ehrler, M., Grassmann, H., Pfenning, P.N., Waha, A., Wohlleber, D., Brockschmidt, F.F., et al. (2012). KIAA1797/FOCAD encodes a novel focal adhesion protein with tumour suppressor function in gliomas. Brain : a journal of neurology 135, 1027–1041.

26. Northcott, P.A., Shih, D.J., Peacock, J., Garzia, L., Morrissy, A.S., Zichner, T., Stutz, A.M., Korshunov, A., Reimand, J., Schumacher, S.E., et al. (2012). Subgroup-specific structural variation across 1,000 medulloblastoma genomes. Nature 488, 49–56.

27. Mardis, E.R. (2014). Sequencing the AML genome, transcriptome, and epigenome. Semin Hematol 51, 250–258.

28. Merker, J.D., Wenger, A.M., Sneddon, T., Grove, M., Zappala, Z., Fresard, L., Waggott, D., Utiramerur, S., Hou, Y., Smith, K.S., et al. (2018). Long-read genome sequencing identifies causal structural variation in a Mendelian disease. Genet Med 20, 159–163.

29. Sedlazeck, F.J., Rescheneder, P., Smolka, M., Fang, H., Nattestad, M., von Haeseler, A., and Schatz, M.C. (2018). Accurate detection of complex structural variations using single-molecule sequencing. Nat Methods 15, 461–468.

30. Cretu Stancu, M., van Roosmalen, M.J., Renkens, I., Nieboer, M.M., Middelkamp, S., de Ligt, J., Pregno, G., Giachino, D., Mandrile, G., Espejo Valle-Inclan, J., et al. (2017). Mapping and phasing of structural variation in patient genomes using nanopore sequencing. Nat Commun 8, 1326.

31. De Coster, W., De Rijk, P., De Roeck, A., De Pooter, T., D’Hert, S., Strazisar, M., Sleegers, K., and Van Broeckhoven, C. (2019). Structural variants identified by Oxford Nanopore PromethION sequencing of the human genome. Genome Res 29, 1178–1187.

32. Kloosterman, W.P., Hoogstraat, M., Paling, O., Tavakoli-Yaraki, M., Renkens, I., Vermaat, J.S., van Roosmalen, M.J., van Lieshout, S., Nijman, I.J., Roessingh, W., et al. (2011). Chromothripsis is a common mechanism driving genomic rearrangements in primary and metastatic colorectal cancer. Genome biology 12, R103.

33. Qian, M., Zhang, H., Kham, S.K., Liu, S., Jiang, C., Zhao, X., Lu, Y., Goodings, C., Lin, T.N., Zhang, R., et al. (2017). Whole-transcriptome sequencing identifies a distinct subtype of acute lymphoblastic leukemia with predominant genomic abnormalities of EP300 and CREBBP. Genome Res 27, 185–195.

34. Rosati, E., Baldoni, S., De Falco, F., Del Papa, B., Dorillo, E., Rompietti, C., Albi, E., Falzetti, F., Di Ianni, M., and Sportoletti, P. (2018). NOTCH1 Aberrations in Chronic Lymphocytic Leukemia. Front Oncol 8, 229.

35. Aljedai, A., Buckle, A.M., Hiwarkar, P., and Syed, F. (2015). Potential role of Notch signalling in CD34+ chronic myeloid leukaemia cells: cross-talk between Notch and BCR-ABL. PLoS One 10, e0123016.

36. Engels, H., Ehrbrecht, A., Zahn, S., Bosse, K., Vrolijk, H., White, S., Kalscheuer, V., Hoovers, J.M., Schwanitz, G., Propping, P., et al. (2003). Comprehensive analysis of human subtelomeres with combined binary ratio labelling fluorescence in situ hybridisation. Eur J Hum Genet 11, 643–651.

37. Genomics, B. (2019). Bionano Solve Theory of Operation: Bionano EnFocusTM FSHD Analysis. Document Number: 30321, Document Revision: A.

38. Luebeck, J., Coruh, C., Dehkordi, S.R., Lange, J.T., Turner, K.M., Deshpande, V., Pai, D.A., Zhang, C., Rajkumar, U., Law, J.A., et al. (2020). AmpliconReconstructor: Integrated analysis of NGS and optical mapping resolves the complex structures of focal amplifications in cancer. bioRxiv, 2020.2001.2022.916031.

39. Sharim, H., Grunwald, A., Gabrieli, T., Michaeli, Y., Margalit, S., Torchinsky, D., Arielly, R., Nifker, G., Juhasz, M., Gularek, F., et al. (2019). Long-read single-molecule maps of the functional methylome. Genome Res 29, 646–656.

40. Ligtenberg, M.J., Kuiper, R.P., Chan, T.L., Goossens, M., Hebeda, K.M., Voorendt, M., Lee, T.Y., Bodmer, D., Hoenselaar, E., Hendriks-Cornelissen, S.J., et al. (2009). Heritable somatic methylation and inactivation of MSH2 in families with Lynch syndrome due to deletion of the 3’ exons of TACSTD1. Nat Genet 41, 112–117.

