## Supplementary Figures 1-4 for "Next generation cytogenetics: comprehensive assessment of 48 leukemia genomes by genome imaging"

### Suppl. Figure 1. Examples of filter adjustment for CNV calling

**a)** *5q22.1qter loss: high-confidence CNV-calling (top) and low confidence (bottom)*

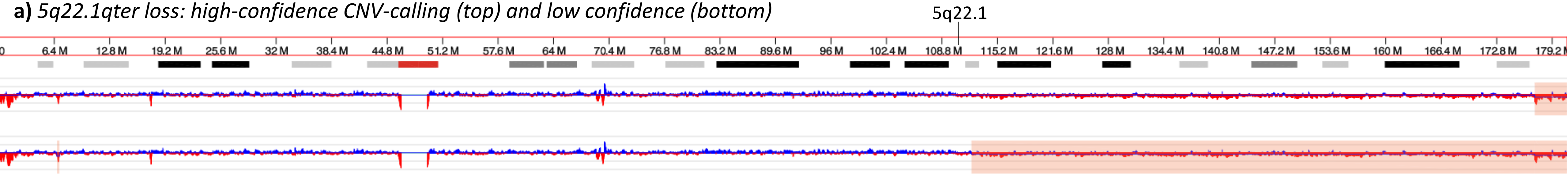


**b)** *Monosomy 18: high-confidence CNV-calling (top) and low confidence (bottom)*

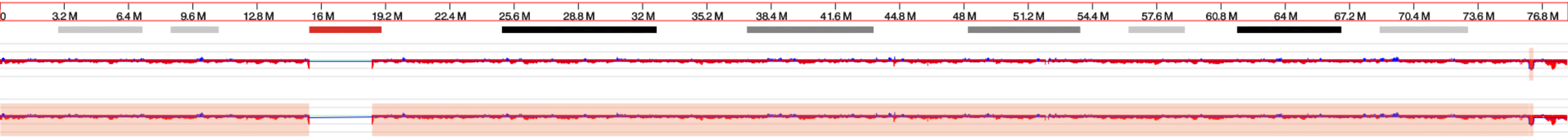


**c)** *5q21.3q34 loss with varying VAFs: high confidence CNV-calling (top) and low confidence (bottom)*

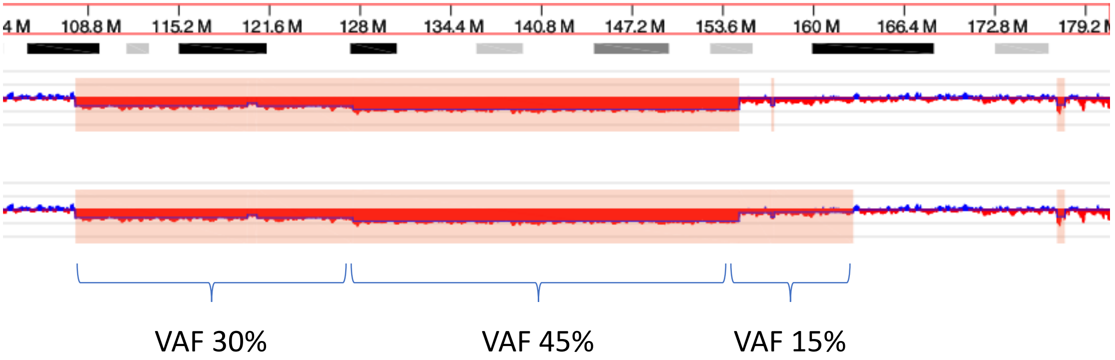

**Suppl. Figure 2: Circos plots of all complex cases**

**Left row shows circos plots with high stringency filters for SVs and CNVs, middle row shows circos plots with low stringency filters for SVs (translocations only) and right row shows circos plots with low stringency filters for CNV calling.**

Sample 38

#### high-confidence

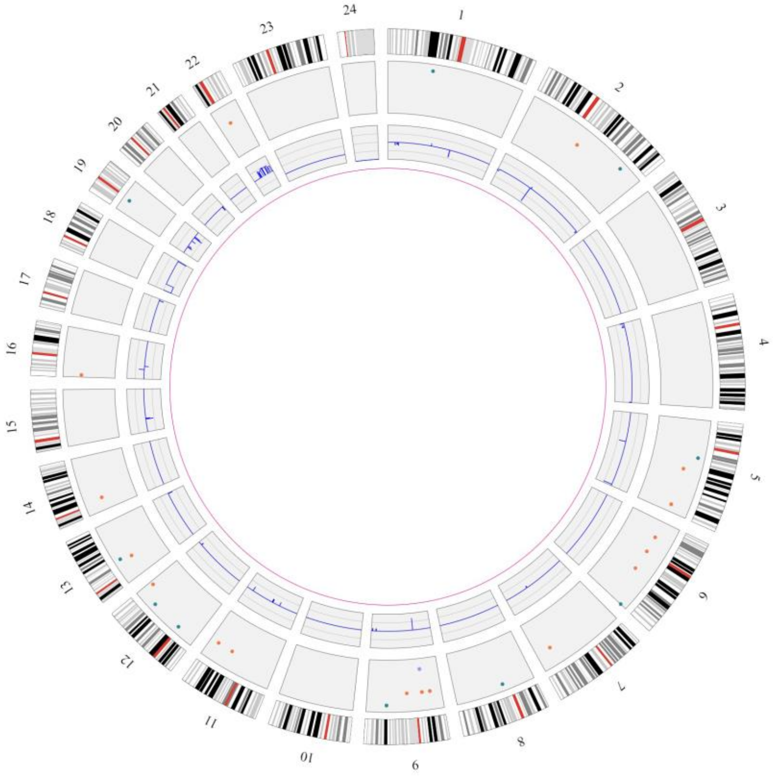

#### Lower confidence translocations

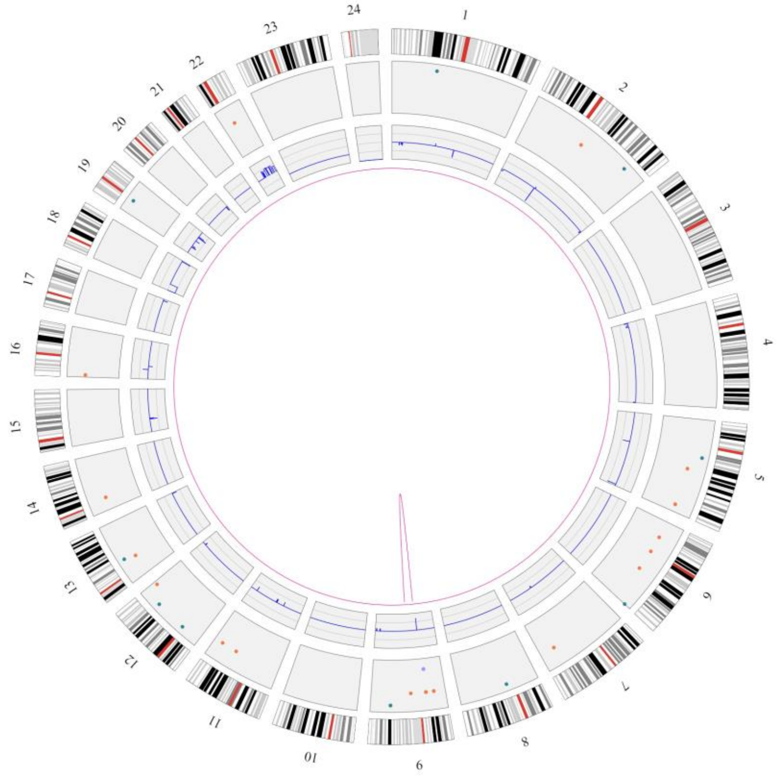

#### Lower confidence CNVs

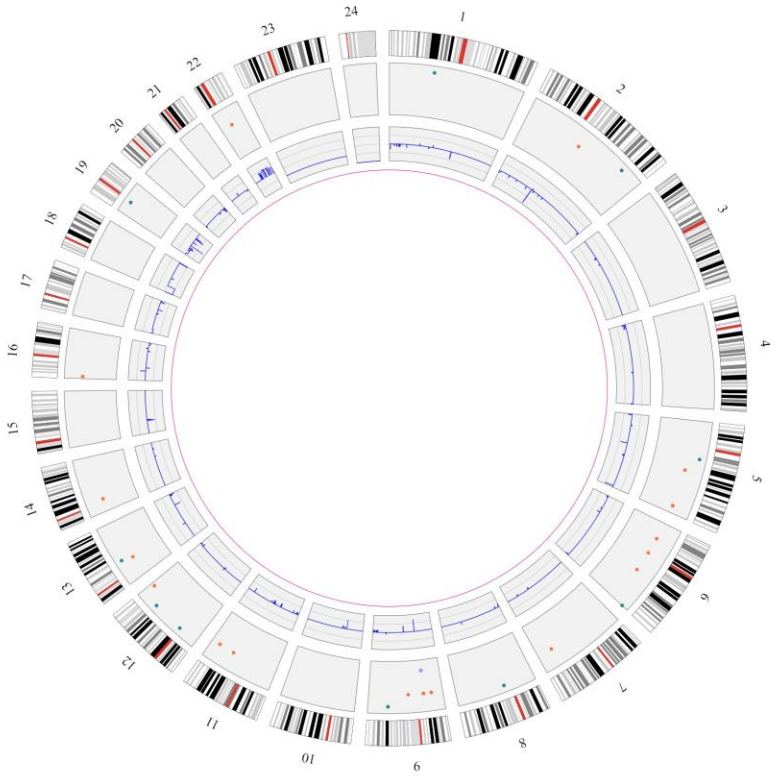

### Sample 39

#### high-confidence

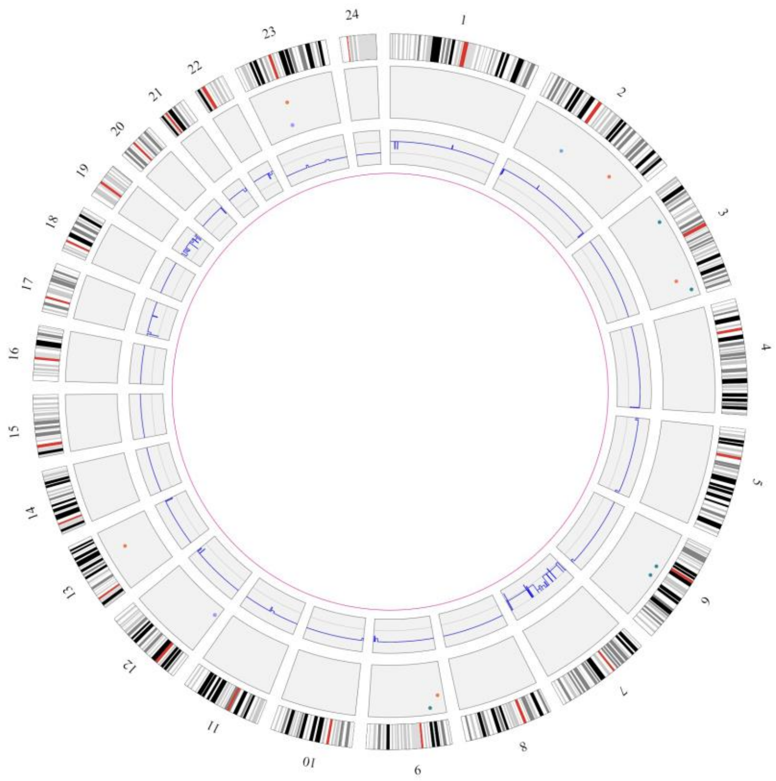


#### Lower confidence translocations

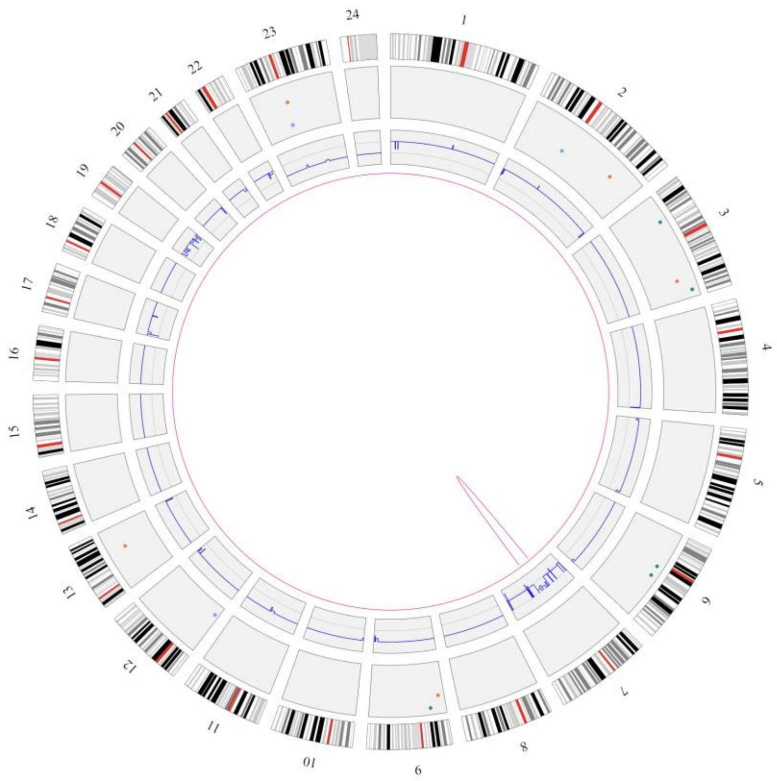


#### Lower confidence CNVs

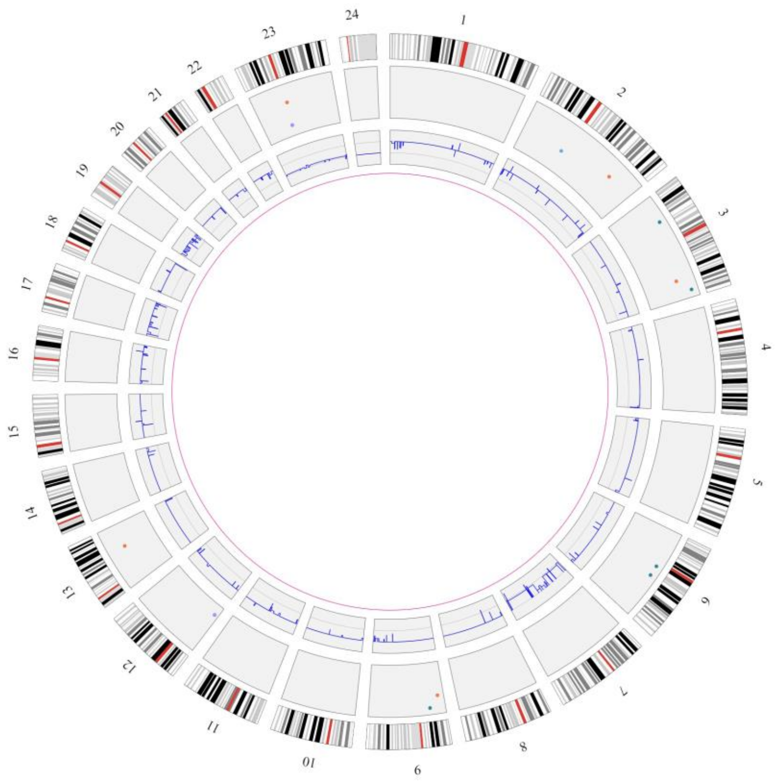

Sample 40

high-confidence

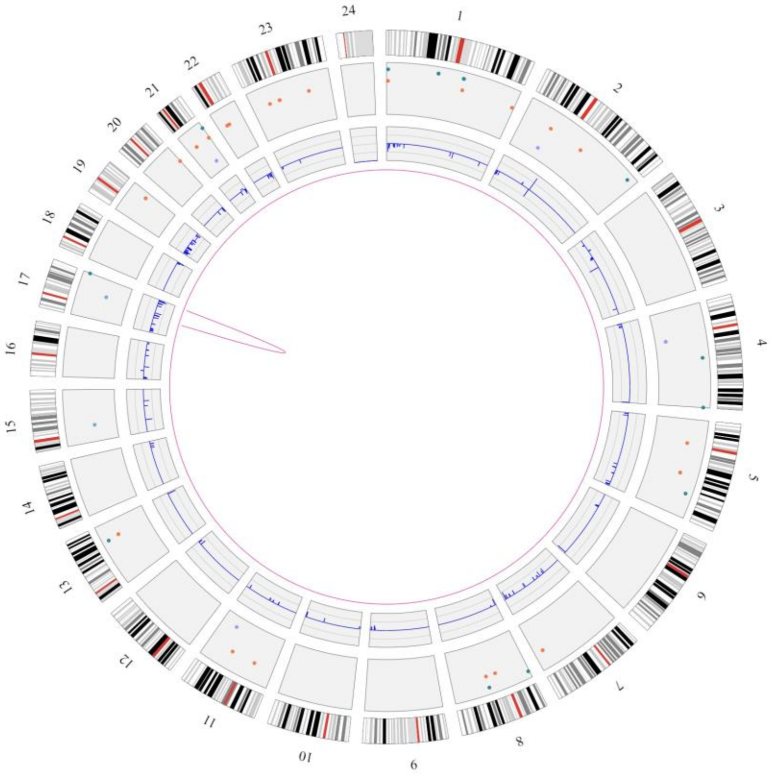


Lower confidence translocations

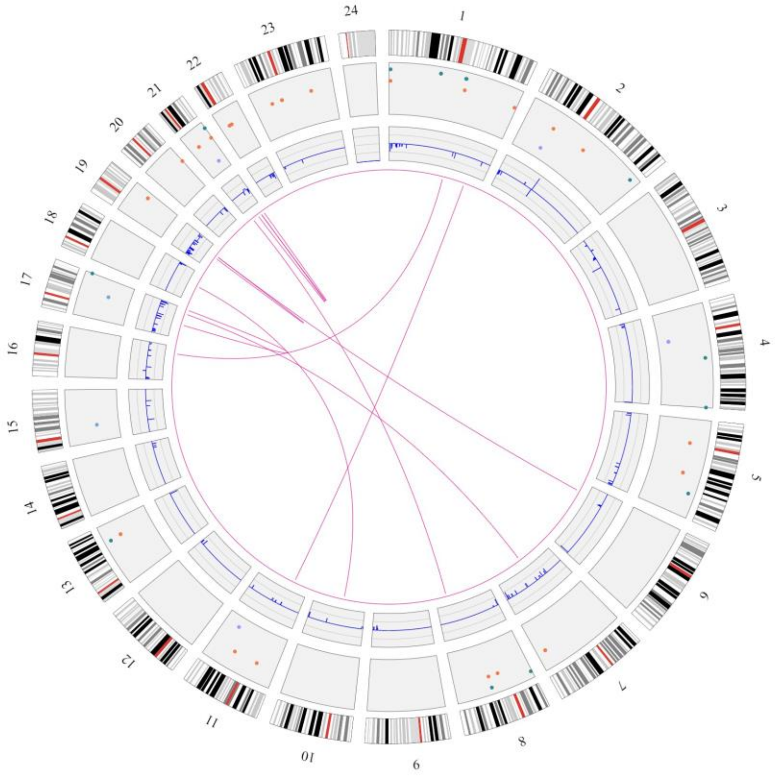


Lower confidence CNVs

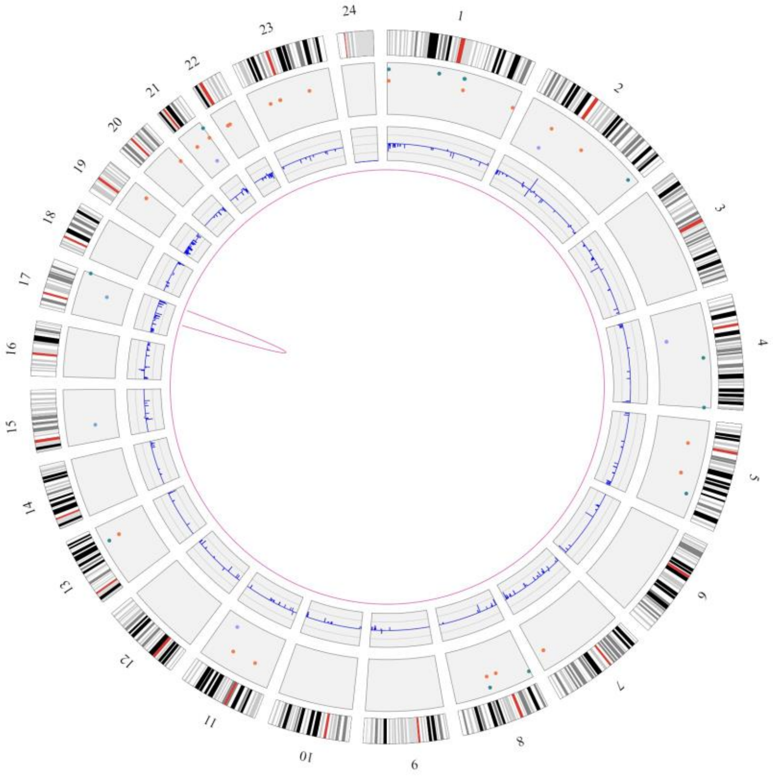

Sample 41

#### high-confidence

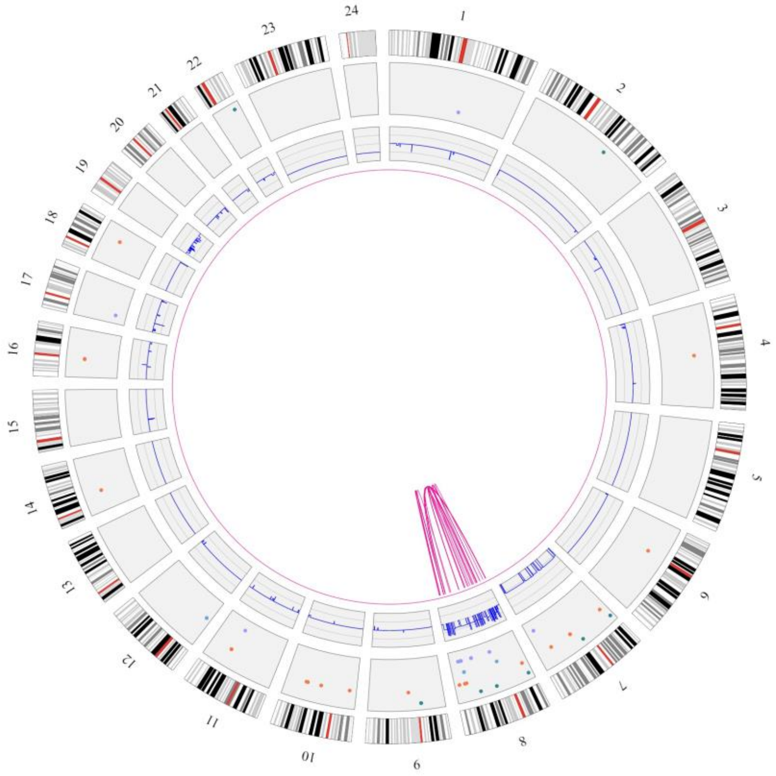


#### Lower confidence translocations

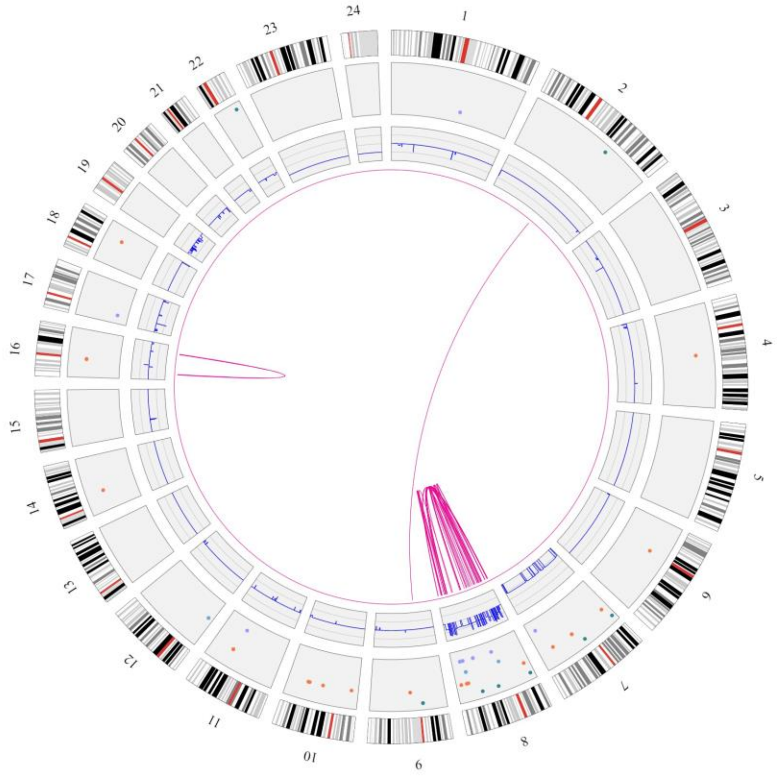


#### Lower confidence CNVs

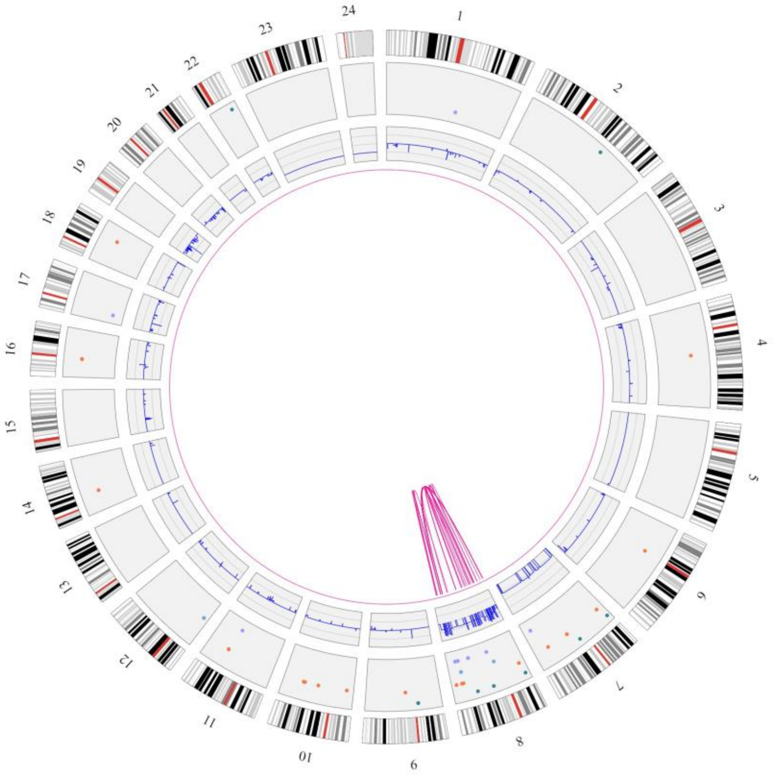

Sample 42

#### high-confidence

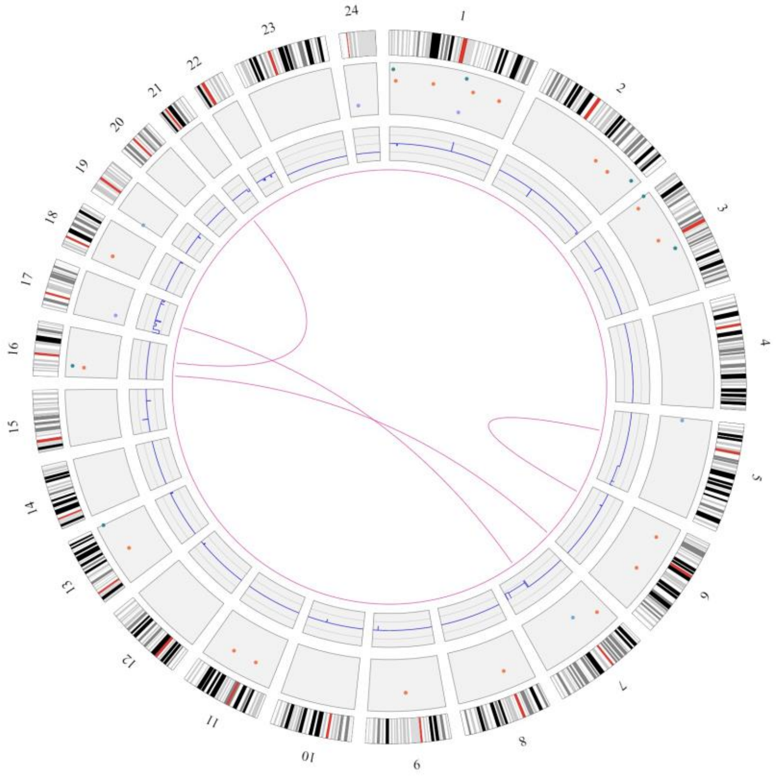

#### Lower confidence translocations

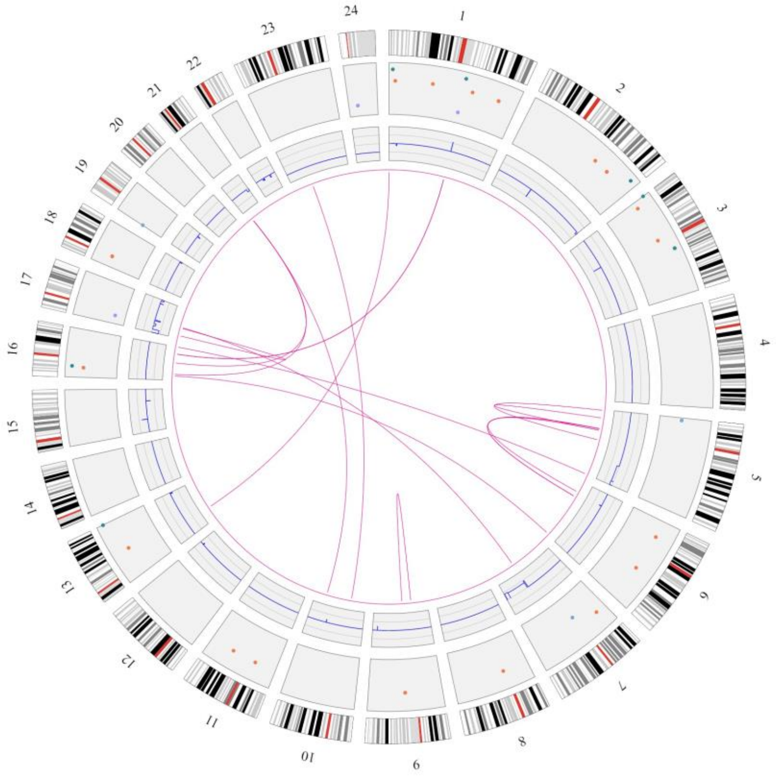

#### Lower confidence CNVs

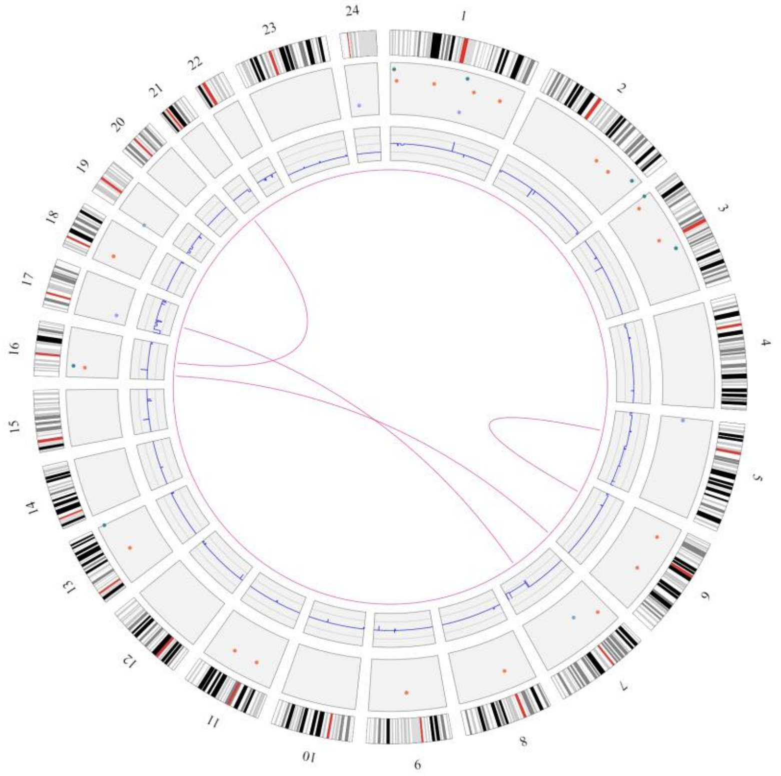

Sample 43

#### high-confidence

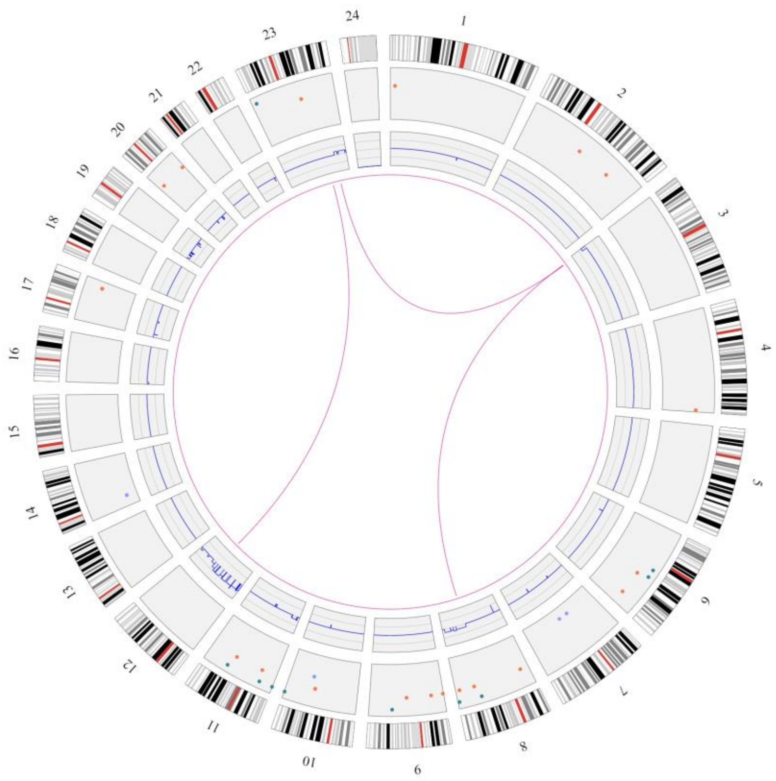


#### Lower confidence translocations

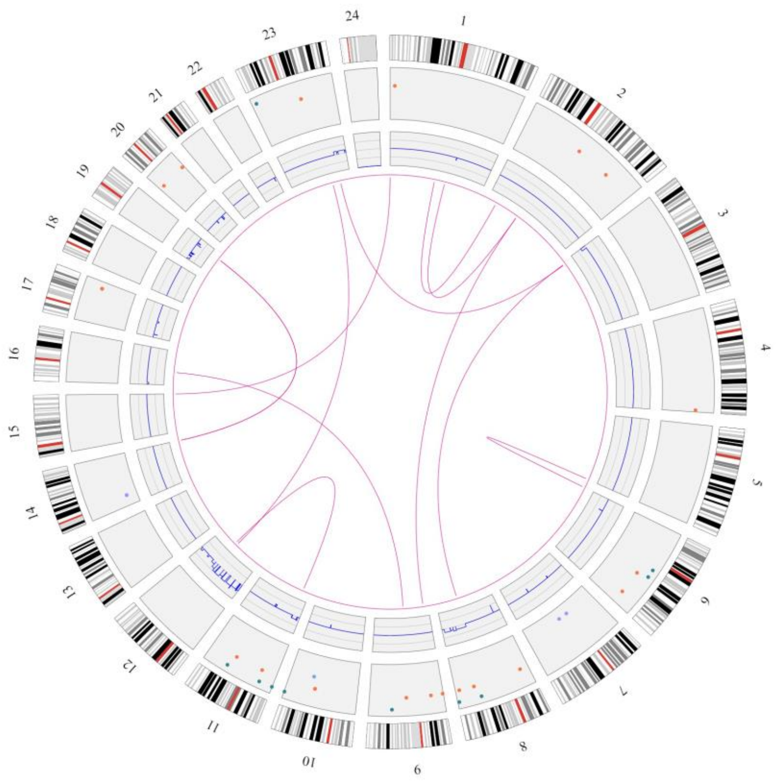


#### Lower confidence CNVs

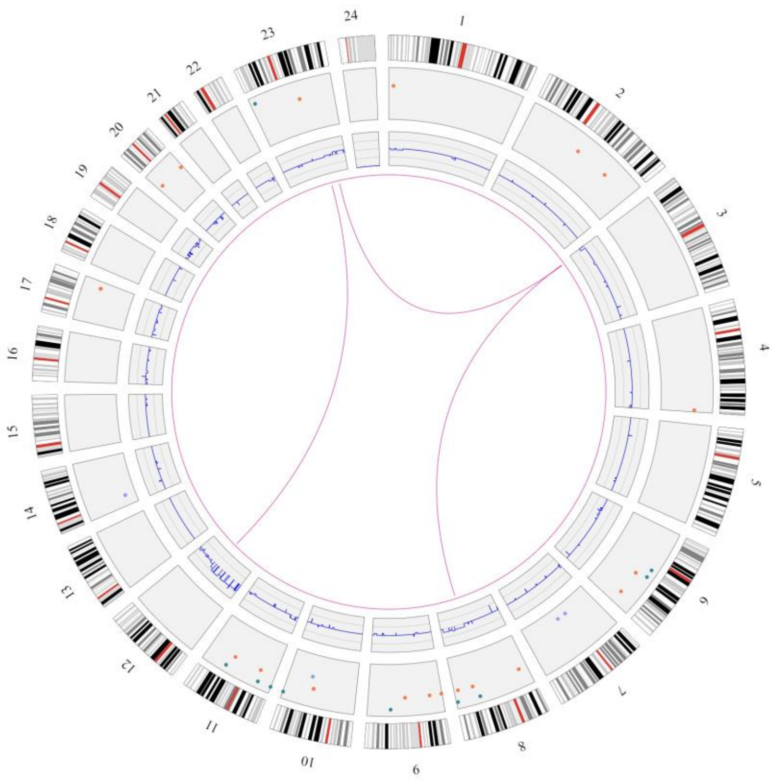

### Sample 44

#### high-confidence

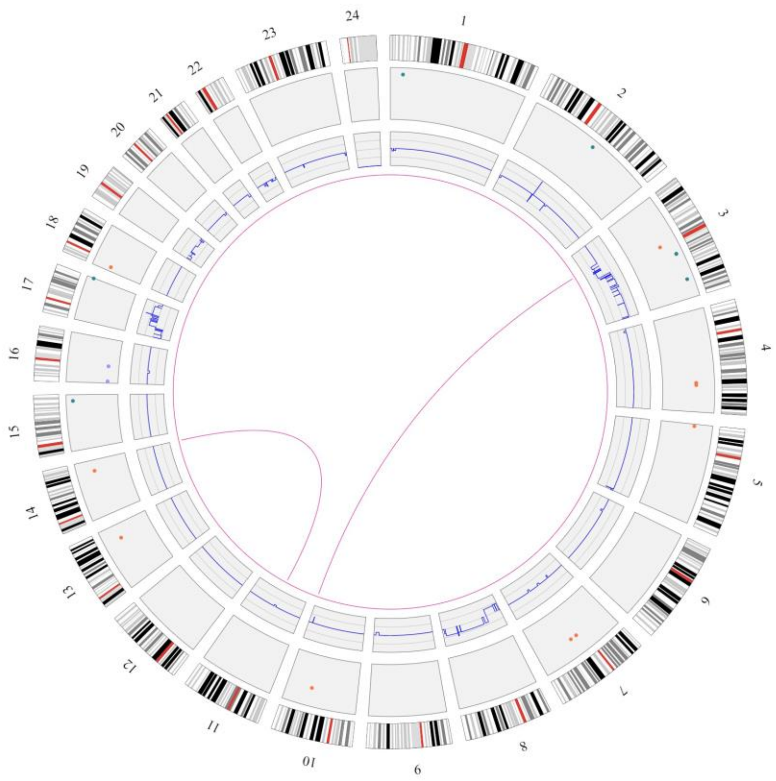


#### Lower confidence translocations

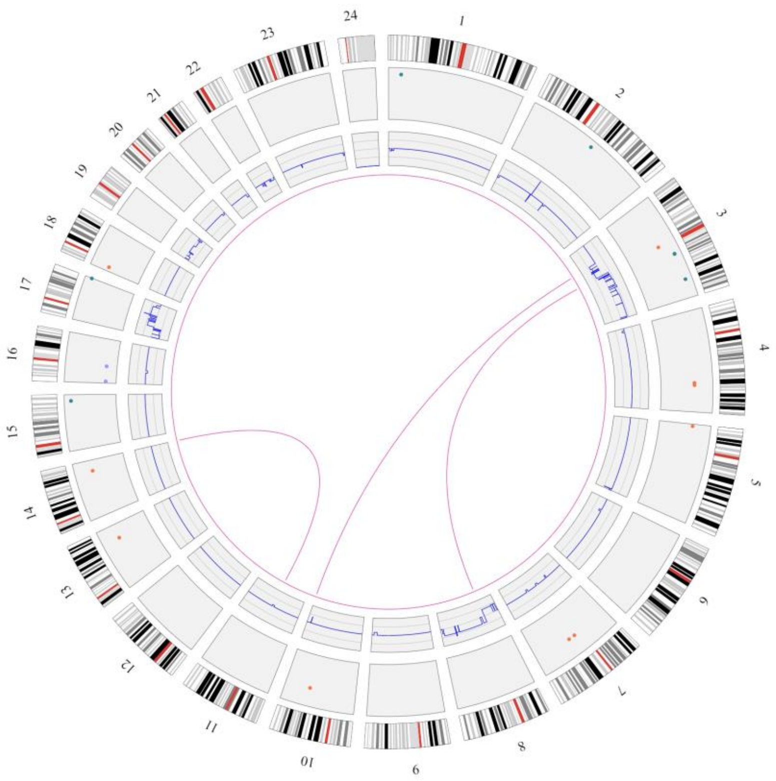


#### Lower confidence CNVs

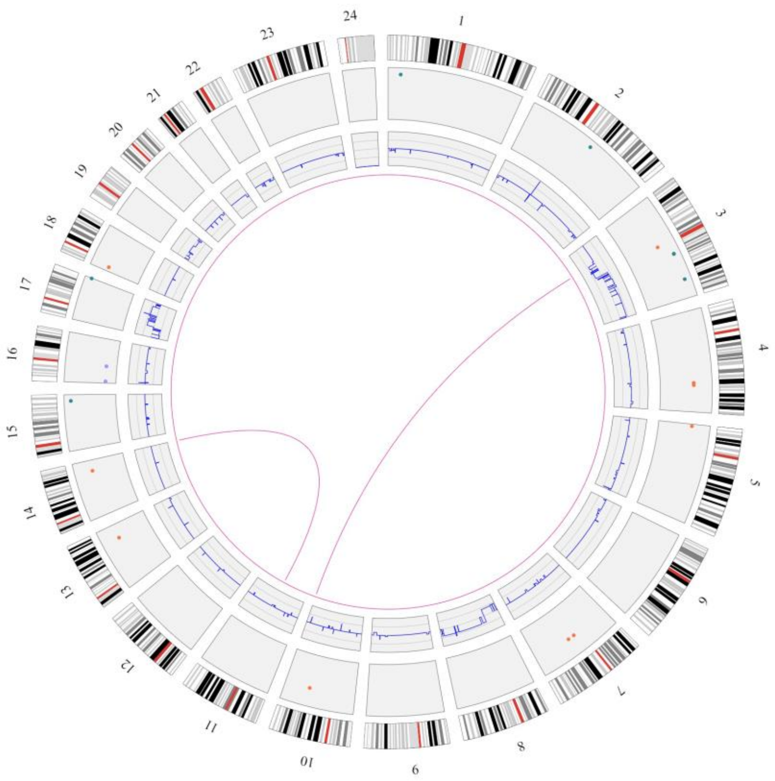

Sample 45

high-confidence

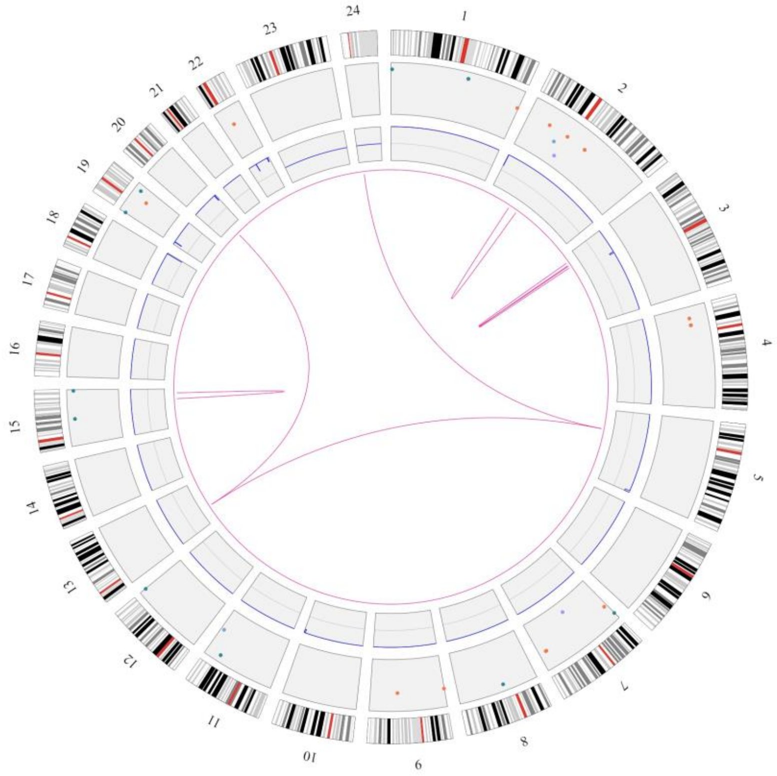


Lower confidence translocations

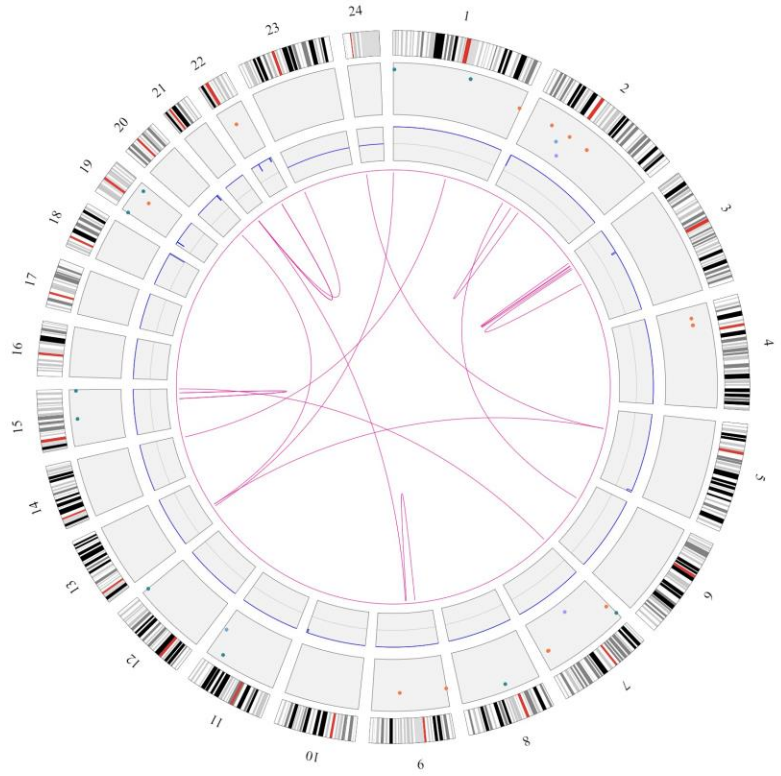


Lower confidence CNVs

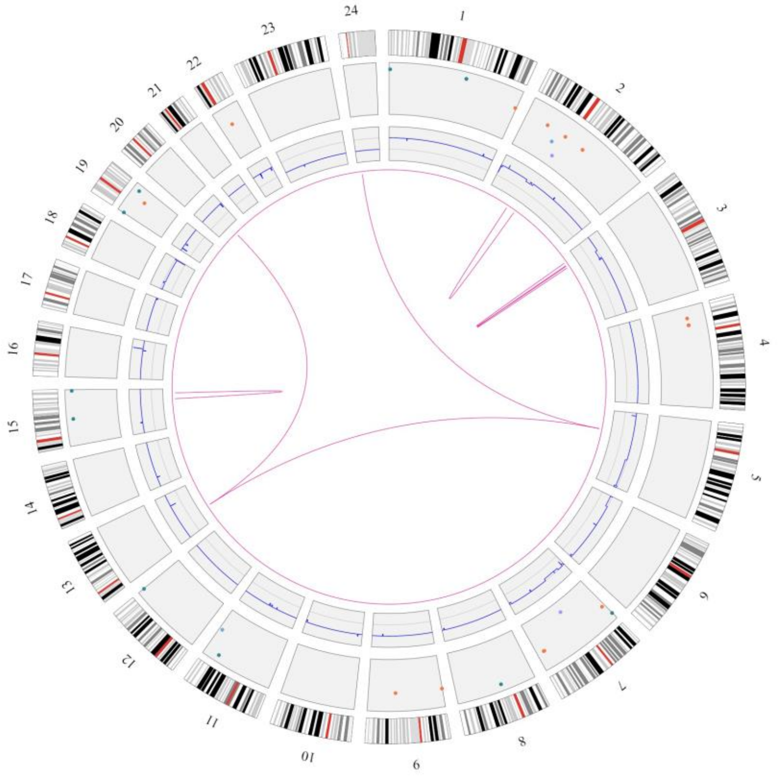

Sample 46

high-confidence

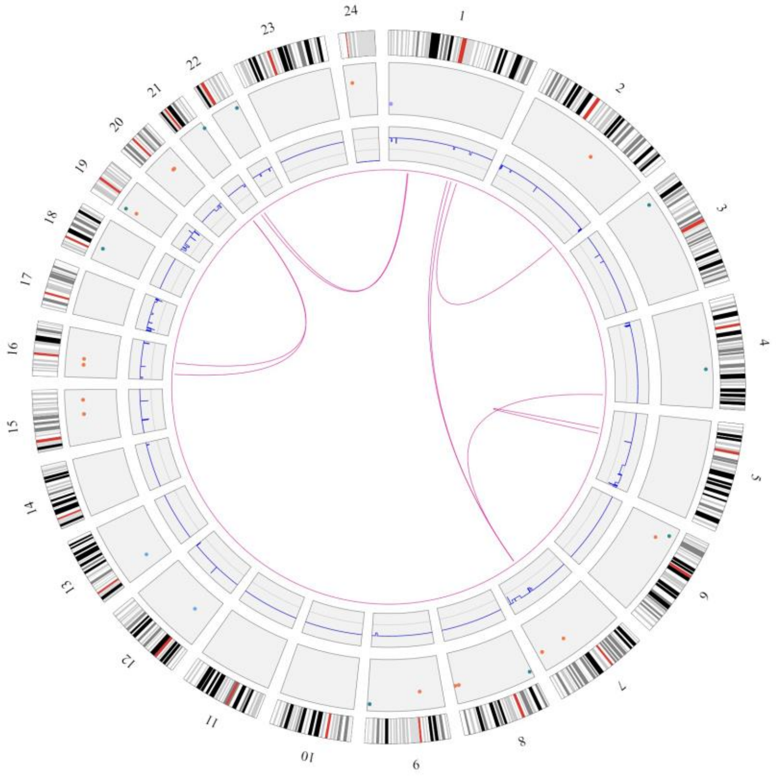


Lower confidence translocations

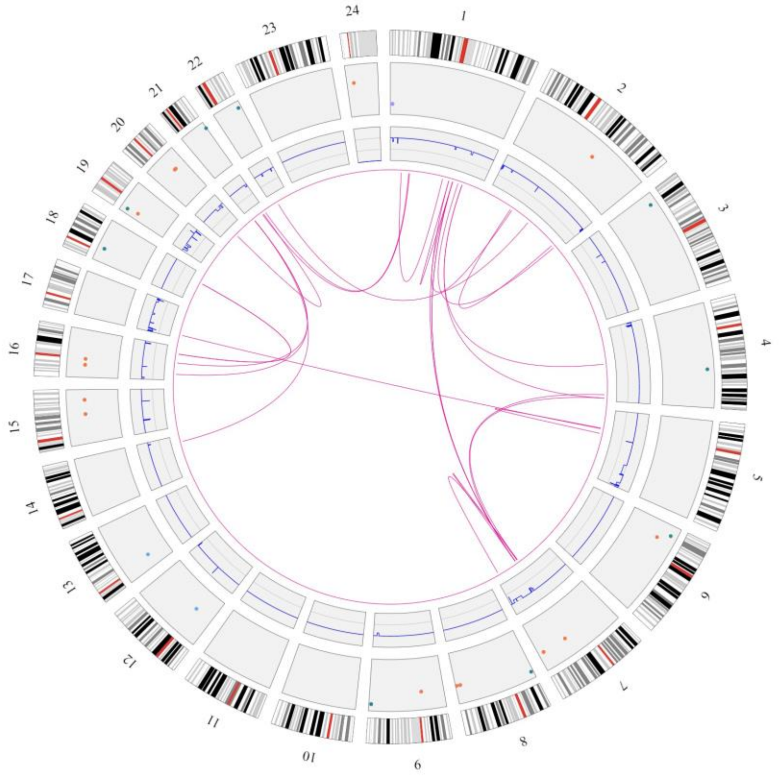


Lower confidence CNVs

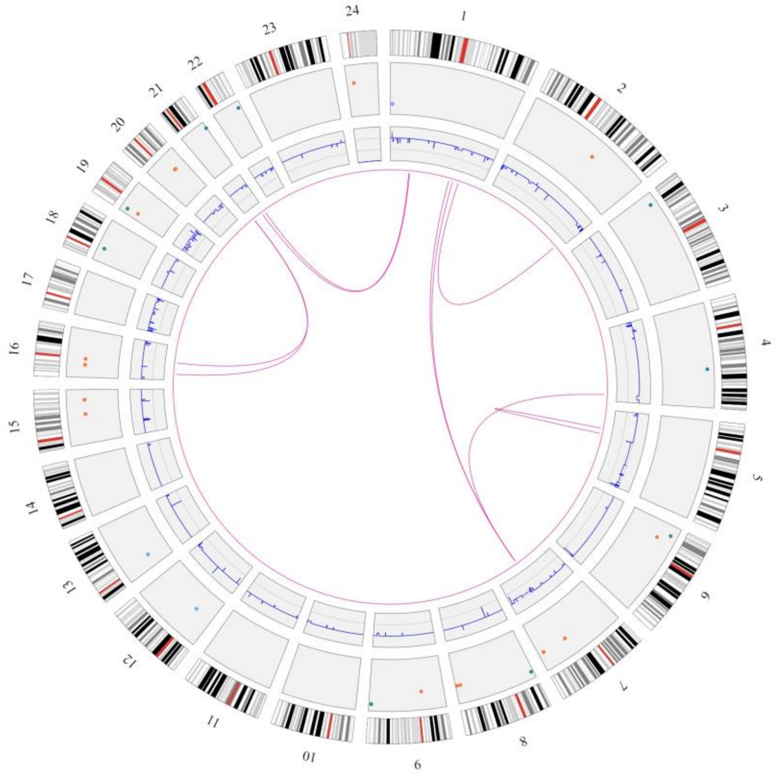

### Sample 47

#### high-confidence

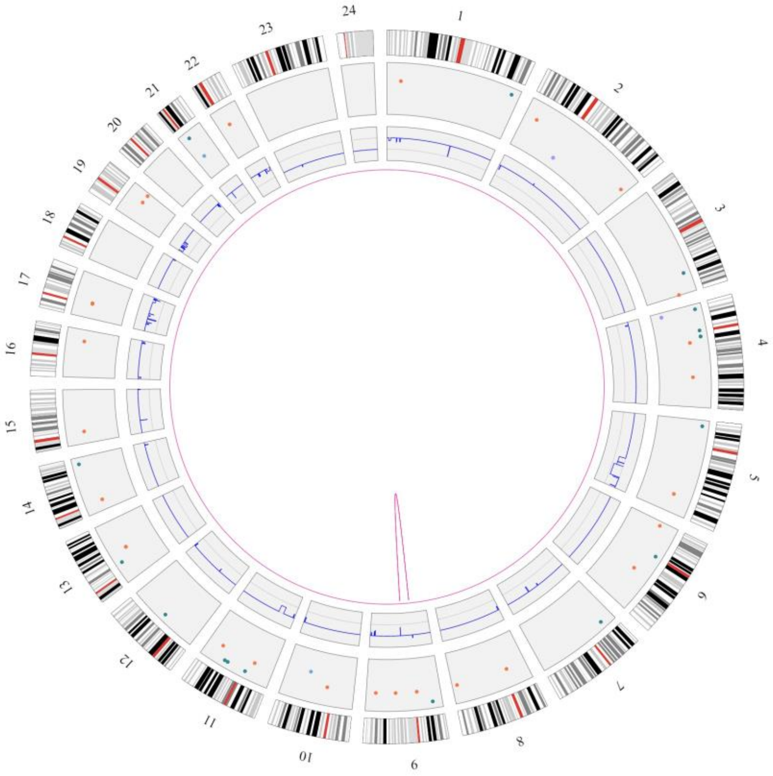


#### Lower confidence translocations

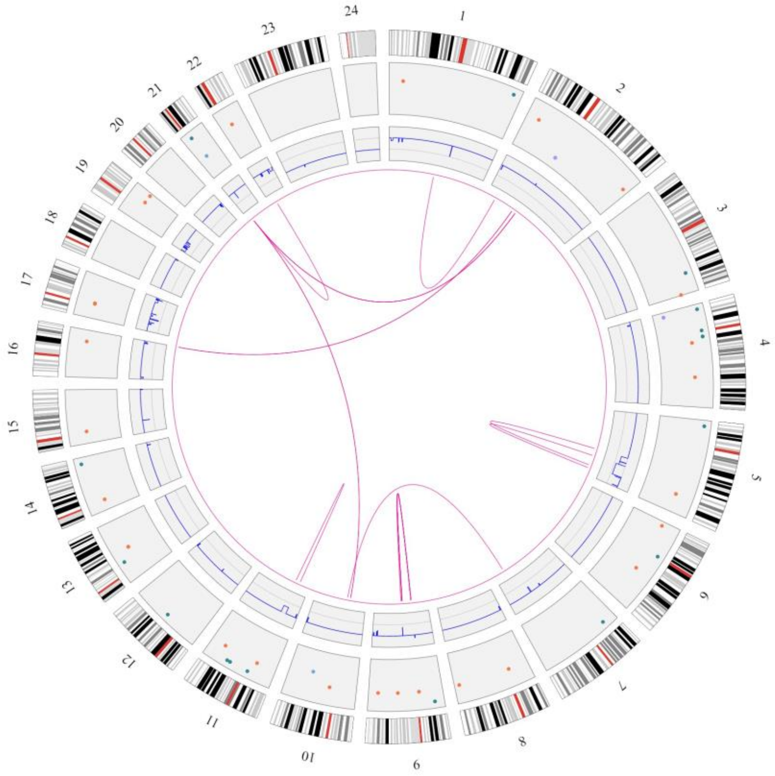


#### Lower confidence CNVs

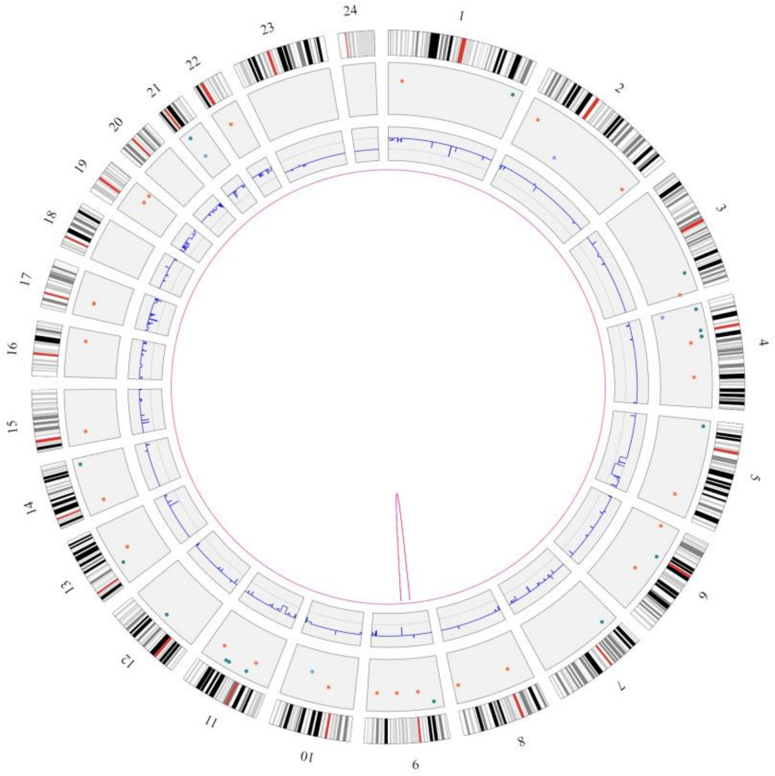

Sample 48

high-confidence

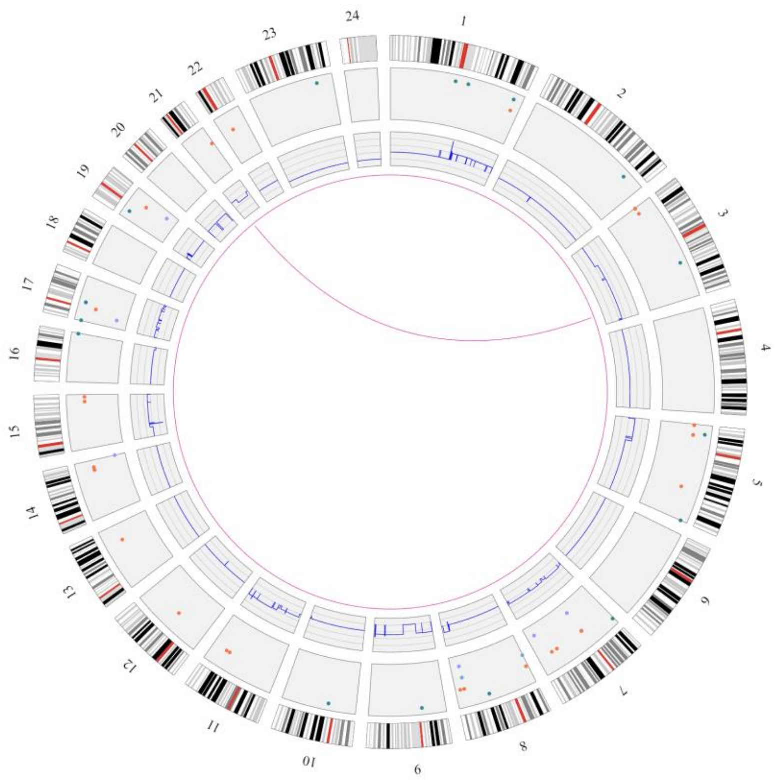


Lower confidence translocations

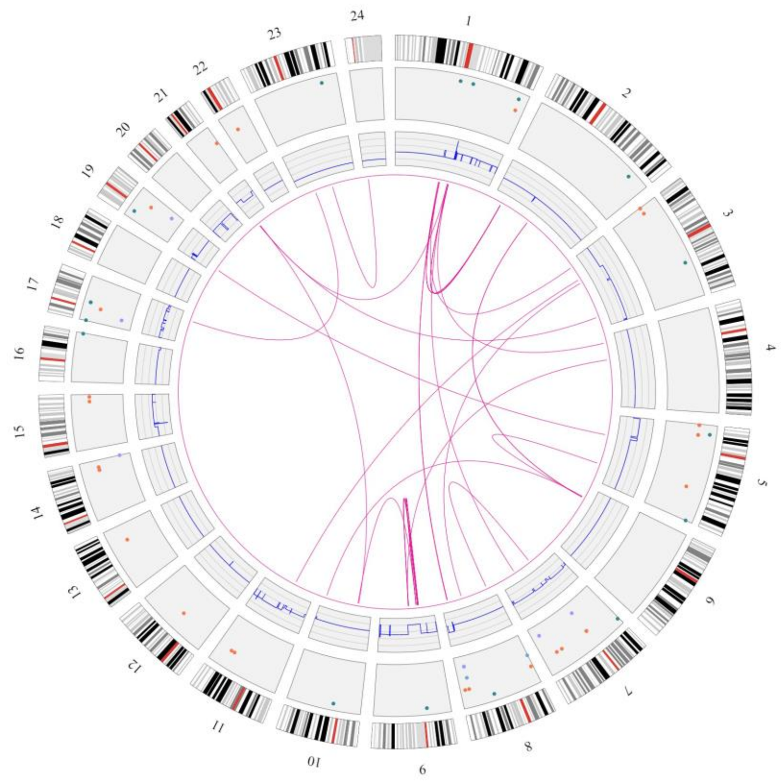


Lower confidence CNVs

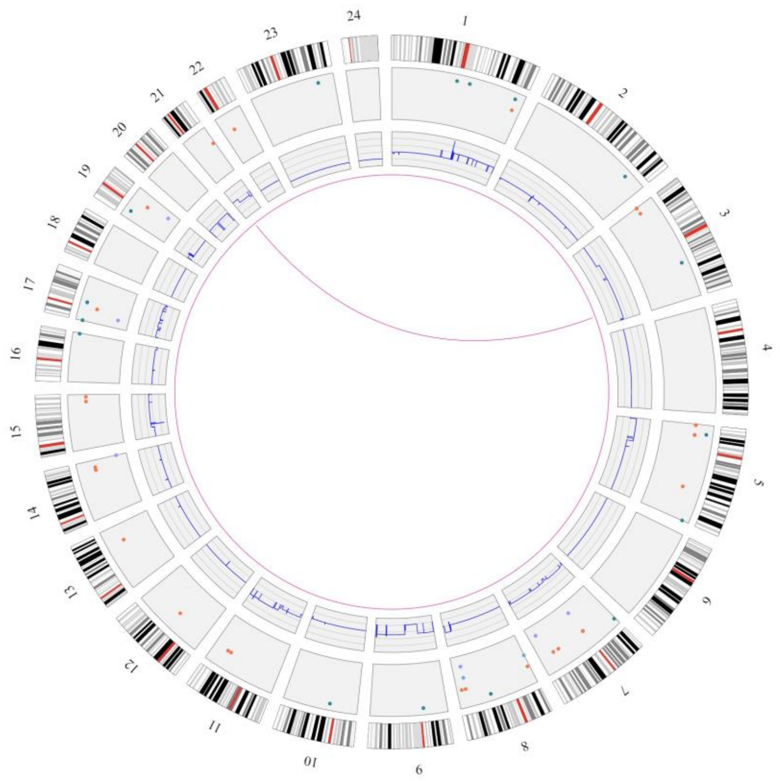

#### Suppl. Figure 3. Fusion gene detection

**a)** Sample 46

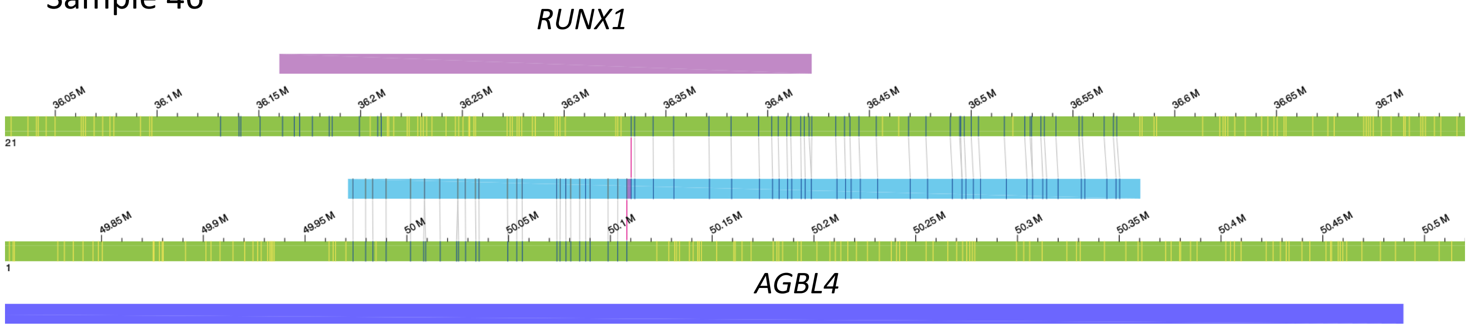


**b)** Sample 14

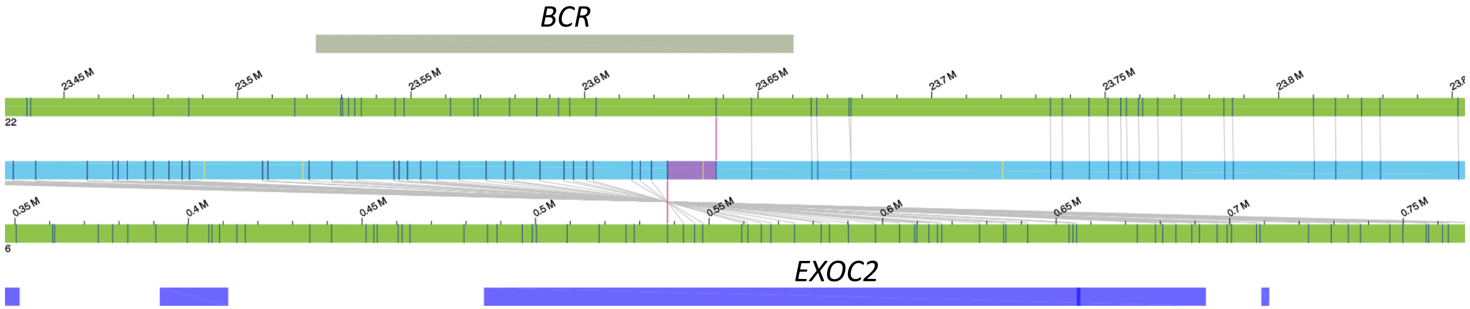

**Suppl. Figure 4: Comparison of CNV-microarray data and genome imaging.**

a)

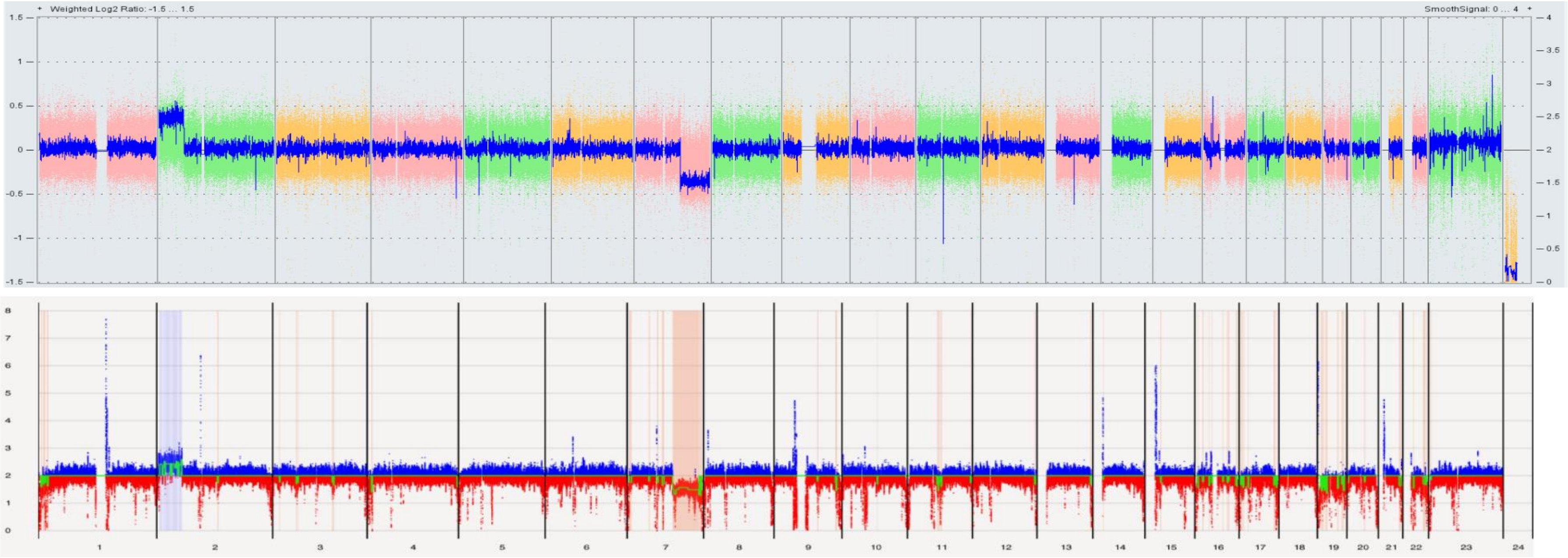


Sample 2:
unbalanced
t(2;7)(p16.3;q21.3)

b)

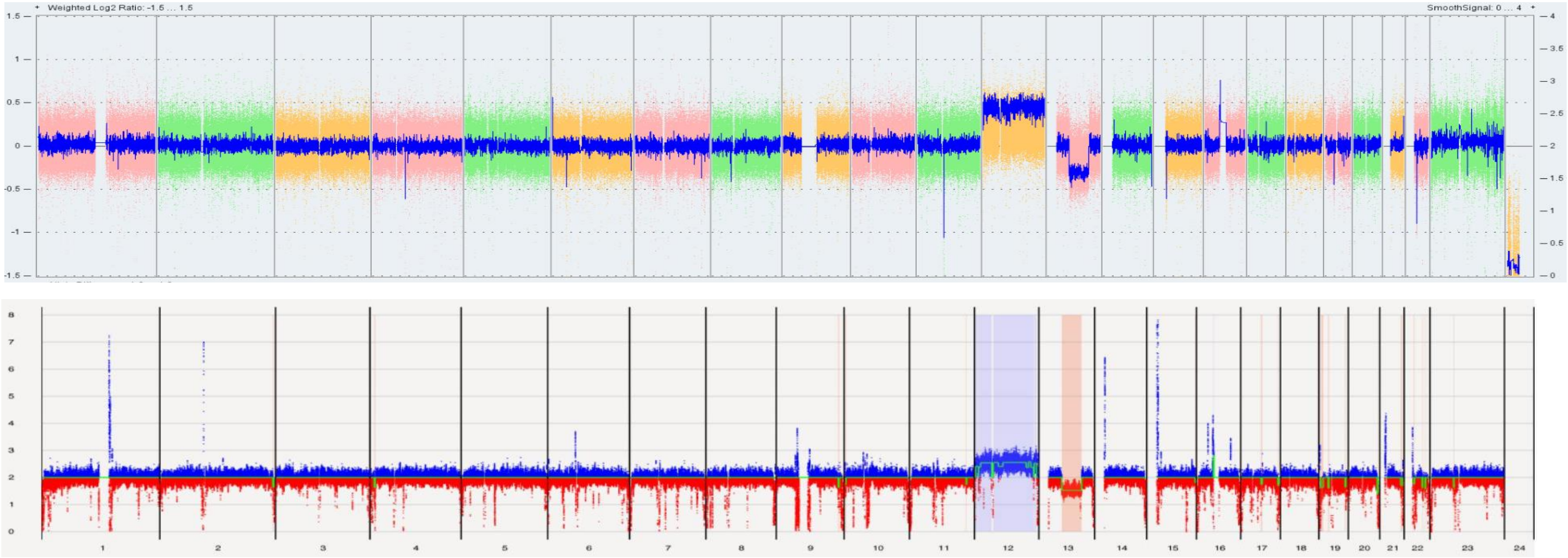


Sample 9
Trisomy 12 and
del13q14.13q31.2
